# Cooperative host-microbe metabolism of a plant toxin in bees

**DOI:** 10.1101/2022.08.25.505265

**Authors:** Erick V. S. Motta, Alejandra Gage, Thomas E. Smith, Kristin J. Blake, Waldan K. Kwong, Ian M. Riddington, Nancy A. Moran

**Author notes:** Corresponding authors: Erick V. S. Motta and Nancy A. Moran 2506 Speedway, Austin, TX 78712, and.

## Abstract

During pollination, bees are exposed to a myriad of xenobiotics, including plant metabolites, which may exert a wide range of effects on their health. Although bees encode enzymes that help in the metabolism of xenobiotics, they still have reduced detoxification gene diversity when compared to other insects, and may rely on other components of their physiology, such as the microbiota, to degrade potentially toxic molecules. In this study, we show that amygdalin, a cyanogenic glycoside found in honey bee-pollinated almond trees, can be metabolized by both bees and members of the gut microbiota. In microbiota-deprived bees, amygdalin is degraded into prunasin, leading to prunasin accumulation in the midgut and hindgut. In microbiota-colonized bees, on the other hand, amygdalin is degraded even further, and prunasin does not accumulate in the gut, suggesting that the microbiota contribute to the full degradation of amygdalin into hydrogen cyanide. *In vitro* experiments demonstrated that amygdalin degradation by bee gut bacteria is strain-specific and not characteristic of a particular genus or species. We found strains of *Bifidobacterium*, *Bombilactobacillus* and *Gilliamella* that can degrade amygdalin, and the degradation mechanism appears to vary since only some strains produce prunasin as an intermediate. Finally, we investigated the basis of degradation in *Bifidobacterium* wkB204, a strain that fully degrades amygdalin. We found overexpression and secretion of several carbohydrate-degrading enzymes, including one in glycoside hydrolase family 3 (GH3). We expressed this GH3 in *Escherichia coli* and detected prunasin as a byproduct when cell lysates were cultured with amygdalin, supporting its contribution to amygdalin degradation. These findings demonstrate that both host and microbiota can act together to metabolize dietary plant metabolites. How amygdalin degradation into hydrogen cyanide affects bee health remains to be elucidated.

## Introduction

During pollination, bees are rewarded with energy-rich nectar, a solution of sugars composed mainly of glucose, fructose, and sucrose, and with pollen, their primary source of proteins, lipids, and vitamins. Nectar and pollen rewards, however, may also contain other sugars and secondary metabolites produced by plants as defenses against infections and herbivores (1). As generalist pollinators, bees are exposed to a wide range of plant metabolites (2). Even when in low concentrations, these metabolites can have a range of effects on bee behavior and health, from negative to neutral to positive, and can be involved in attraction or deterrence (3–5). Interestingly, some bee species cannot detect naturally occurring concentrations of certain nectar metabolites, such as quinine, nicotine, caffeine and amygdalin (6). This poor acuity may lead to long-term side effects depending on the toxicity of the metabolite.

A plant secondary metabolite to which bees may be chronically exposed is amygdalin, a cyanogenic glycoside found in the nectar and pollen of crops, such as almonds, apples, cherries, and nectarines, for which the western honey bee, *Apis mellifera* is the primary pollinator (7). Amygdalin must be degraded to exert a toxic effect (8). Degradation occurs during tissue damage, such as chewing by herbivores, since amygdalin is stored in cell vacuoles and the glycoside hydrolases (GHs) involved in degradation are present in the cytoplasm. During degradation, amygdalin is usually first broken down into prunasin and a glucose molecule. Then, prunasin is broken down into another glucose molecule and mandelonitrile, with the latter compound converted into benzaldehyde and hydrogen cyanide. Hydrogen cyanide is a toxic molecule that leads to acute poisoning in animals (9–14) as it interferes with the electron transport chain during oxidative phosphorylation (15). Interestingly, some bees are not deterred by amygdalin concentrations encountered in almond nectar (up to 15 μM) (7, 16), and can tolerate high concentrations (up to 219 μM) with no effects on survivorship (2, 17). However, exposure to higher doses of amygdalin can lead to malaise symptoms, such as increased time spent upside down and abdomen dragging (18).

Although honey bees and bumble bees have fewer detoxification genes compared to other insects (19, 20), they produce some enzymes that help degrade plant metabolites, such as cytochrome P450 monooxygenases, glutathione transferases, and glycoside hydrolases (GHs) (19, 21, 22). For example, honey bees secrete a GH into their mouths from their hypopharyngeal glands that is then transferred to the midgut where it can potentially catalyze the initial breakdown of glycosides (21, 23), such as the conversion of amygdalin into prunasin. Amygdalin toxicity occurs after ingestion, but not after injection into the hemolymph (18), suggesting that enzymes in the gut achieve the conversion of amygdalin into toxic cyanide. Whether amygdalin is fully processed by bee GHs (21) or by pollen-derived GHs that bees ingest (23) is unknown. Another possibility is that the bee gut microbiota (24–26) contributes to amygdalin degradation, as suggested by the vast arsenal of GHs produced by the dominant bee gut bacterial species (27, 28). Interestingly, amygdalin itself does not show antibacterial effects *in vitro*, and the honey bee gut microbiota appears not to be significantly affected by amygdalin exposure (29). In fact, colony-level exposure to amygdalin may protect bees against some parasites, such as *Lotmaria passim* (29), and reduce the titer of pathogenic viruses (29, 30), suggesting a potential trade-off in metabolizing amygdalin.

Specific members of the bee gut microbiota encode a diverse set of carbohydrate digestive enzymes, including pectin lyases (PLs) and GHs, that help in food processing (27, 31, 32) and detoxification (19, 33). For instance, several *Gilliamella* strains encode PLs and GHs that are involved in the metabolism of pectin and hemicellulose from the pollen cell wall and toxic sugars from nectar or produced during digestion of pectin (31, 32). Some of these sugars, such as mannose, arabinose, xylose and galactose, are indigestible for bees and can cause toxicity if accumulated in the gut (34). Strains of *Bombilactobacillus* and *Lactobacillus* nr. *melliventris* (formerly called *Lactobacillus* Firm-4 and *Lactobacillus* Firm-5, respectively) also contain genes encoding enzymes of mannose metabolism and GHs and potentially contribute to this detoxification mechanism (27, 28, 32, 35). Interestingly, *Bifidobacterium* strains seem to harbor a wider repertoire of GHs than other core members of the bee gut microbiota, but lack PL-related genes (27). Despite the interest in correlating the effects of nectar metabolites with bee behavior and health, there remains a gap in understanding the contributions of the microbiota to degradation of plant metabolites, with only a few studies addressing this issue (27, 31, 36).

In this study, we investigated the contributions of honey bees and their microbiota to amygdalin degradation. We found that breakdown to prunasin can be achieved by hosts without a microbiota and that further degradation can be performed by specific strains of dominant microbiota species. Using biochemical assays, we characterized a GH secreted by bee-associated *Bifidobacterium* strains that can degrade amygdalin and prunasin. These findings shed light on how the combined contributions of host and microbiome enable degradation of a dietary plant metabolite.

## Results

To investigate metabolism by bee gut bacteria, we selected representative strains of four bacterial groups involved in food metabolism in the bee gut: *Bifidobacterium*, *Bombilactobacillus*, *Lactobacillus* nr. *melliventris*, and *Gilliamella* (Figure 1). We cultured these strains in semi-defined media (SDM, Figure 1A) or in nutritionally rich media (MRS or Insectagro, Figure 1B) to assess their susceptibility to amygdalin and their ability to metabolize amygdalin into byproducts, such as prunasin, as analyzed by LC-MS (Figure 1C).

**Figure 1.**
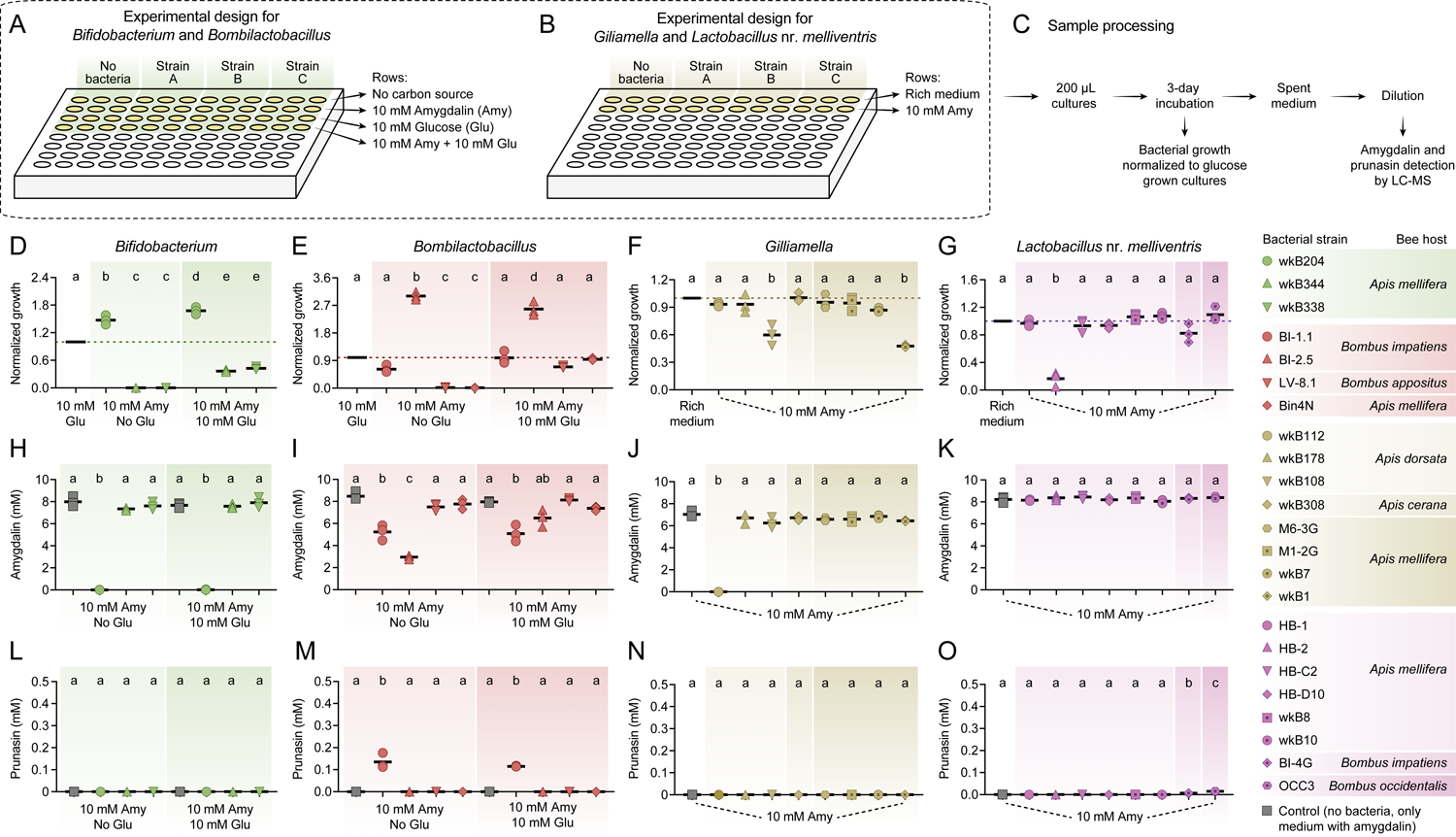
*In vitro* exposure of bee gut bacteria to amygdalin. Experimental design in **(A)** semi-defined or **(B)** nutritionally rich media in 96-well plates. **(C)** Sample processing for LC-MS analysis. **(D)** *Bifidobacterium* and **(E)** *Bombilactobacillus* growth in semi-defined media in the presence of amygdalin (or amygdalin and glucose) normalized to growth in the presence of glucose. **(F)** *Gilliamella* and **(G)** *Lactobacillus* nr. *melliventris* growth in nutritionally rich media in the presence of amygdalin normalized to growth in the absence of amygdalin. Bacterial growth was measured as optical density at 600 nm after 3 days of incubation at 35°C and 5% CO_2_. **(H-K)** Amygdalin and **(L-O)** prunasin concentrations in spent medium of amygdalin (or amygdalin and glucose) grown cultures of *Bifidobacterium*, *Bombilactobacillus*, *Gilliamella* and *Lactobacillus* nr. *melliventris*, respectively. Controls consisted of media with amygdalin (or amygdalin and glucose) but no bacteria. Experiments were performed in three biological replicates. Groups with different letters are significantly different (*P* < 0.01, One-way ANOVA test followed by Tukey’s multiple-comparison test).

### Bee gut bacterial symbionts vary in susceptibility to amygdalin

Strains varied in their ability to cope with different concentrations of amygdalin *in vitro*.

### Bifidobacterium

Three strains isolated from the guts of *Apis mellifera* (wkB204, wkB344 and wkB338) were cultured in the presence of amygdalin and/or glucose as sole carbon sources in SDM (Figure 1A). Only strain wkB204 grew in the presence of amygdalin as the sole carbon source, suggesting that this strain degrades amygdalin and is not susceptible to the potential byproducts (Figure 1D). On the other hand, strains wkB344 and wkB338 grew only when glucose was added, and their growth was hampered if amygdalin was added, indicating a toxic effect on these strains (Figure 1D).

### Bombilactobacillus

Strains isolated from the guts of *Bombus impatiens* (BI-1.1 and BI-2.5), *Bombus appositus* (LV-8.1) and *A. mellifera* (Bin4N) were tested in SDM (Figure 1A). Strains BI-1.1 and BI-2.5 grew in the presence of amygdalin as the sole carbon source, with BI-2.5 growing better than BI-1.1 or the control with glucose as the sole carbon source (Figure 1E). Strains LV-8.1 and Bin4N grew only in the medium with glucose, but their growth was not affected when amygdalin was added (Figure 1E).

### Gilliamella

Strains isolated from the guts of *Apis dorsata* (wkB112, wkB178 and wkB108), *Apis cerana* (wkB308) and *A. mellifera* (M6-3G, M1-2G, wkB7, and wkB1) were cultivated in Insectagro due to the lack of a SDM for these strains (Figure 1B). Most of the strains grew at similar rates in the presence or absence of 10 mM amygdalin, except for wkB108 and wkB1 which exhibited a delay in growth, suggesting susceptibility to amygdalin at the tested concentration (Figure 1F).

### Lactobacillus nr. melliventris

Strains isolated from the guts of *A. mellifera* (HB-1, HB-2, HB-C2, HB-D10, wkB8, and wkB10), *B. impatiens* (BI-4G), and *Bombus occidentalis* (OCC3) were cultivated in rich medium (MRS) since they do not grow well in SDM (Figure 1B). All strains grew in the presence of amygdalin, though HB-2 growth was reduced by adding amygdalin to MRS (Figure 1G).

For most bacterial strains tested, growth was hampered by increasing the concentration of amygdalin from 10 mM to 100 mM (Figure S1A-C). This toxicity is probably related to the presence of amygdalin itself and not to potential byproducts since most strains could not degrade amygdalin. The amygdalin concentrations were chosen to correspond to the glucose concentrations usually added to growth media to investigate carbon source usage by bacteria.

### Specific bee gut bacterial strains degrade amygdalin

Using LC-MS analyses, amygdalin degradation was confirmed for strains that could grow in the presence of amygdalin as the sole carbon source, such as *Bifidobacterium* strain wkB204 (Figure 1H) and *Bombilactobacillus* strains BI-1.1 and BI-2.5 (Figure 1I). Amygdalin was not detected (Figure 1H) or was detected in a lower concentration (Figure 1I and Figure S1D-E) in the spent medium of amygdalin-grown cultures when compared to the initial concentration. In these cases, amygdalin degradation was observed regardless of whether glucose was present. Interestingly, *Bombilactobacillus* strain BI-2.5 prefers to use glucose as carbon source rather than amygdalin when both compounds are present in the medium (Figure 1I). For *Bifidobacterium* strain wkB204 and *Bombilactobacillus* strain BI-1.1, on the other hand, similar levels of amygdalin degradation were detected in cultures with or without glucose (Figure 1H-I).

*Gilliamella* and *Lactobacillus* nr. *melliventris* strains were cultivated in nutritionally rich media, and therefore amygdalin degradation was primarily investigated by LC-MS of spent medium. We observed amygdalin degradation only for *Gilliamella* strain wkB112 (Figure 1J and Figure S1F). The use of nutritionally rich media for these strains may have masked the ability of some strains to degrade amygdalin, as they had glucose as an alternative carbon source (Figure 1J-K).

### Different mechanisms of amygdalin degradation by bee gut bacteria

Metabolism of amygdalin by *Bifidobacterium* strain wkB204 and *Bombilactobacillus* strain BI-1.1 produces prunasin as a byproduct (Figure 1L-M and Figure S1G-H), although prunasin was only detected in wkB204 cultures after providing an excessive amount of amygdalin (Figure S1G). This suggests that wkB204 and BI-1.1 encode enzymes to break down the glycosidic bond between the glucose residues in the amygdalin structure, releasing prunasin and one glucose molecule, which can then be used as carbon source by these bacteria. On the other hand, prunasin was not produced by *Bombilactobacillus* strain BI-2.5 or *Gilliamella* strain wkB112 (Figure 1M-N) even after adding excess amygdalin (Figure S1H-I). Therefore, BI-2.5 and wkB112 seem to metabolize amygdalin in a different way than wkB204 and BI-1.1, probably by breaking down the glycosidic bond that links the two glucose residues to the aglycone, releasing a disaccharide and mandelonitrile into the medium. These mechanisms are corroborated by LC-MS analyses of spent medium taken from these cultures on a daily census (Figure 2). These results suggest that amygdalin breakdown via a prunasin intermediate is limited to wkB204 and BI-1.1.

**Figure 2.**
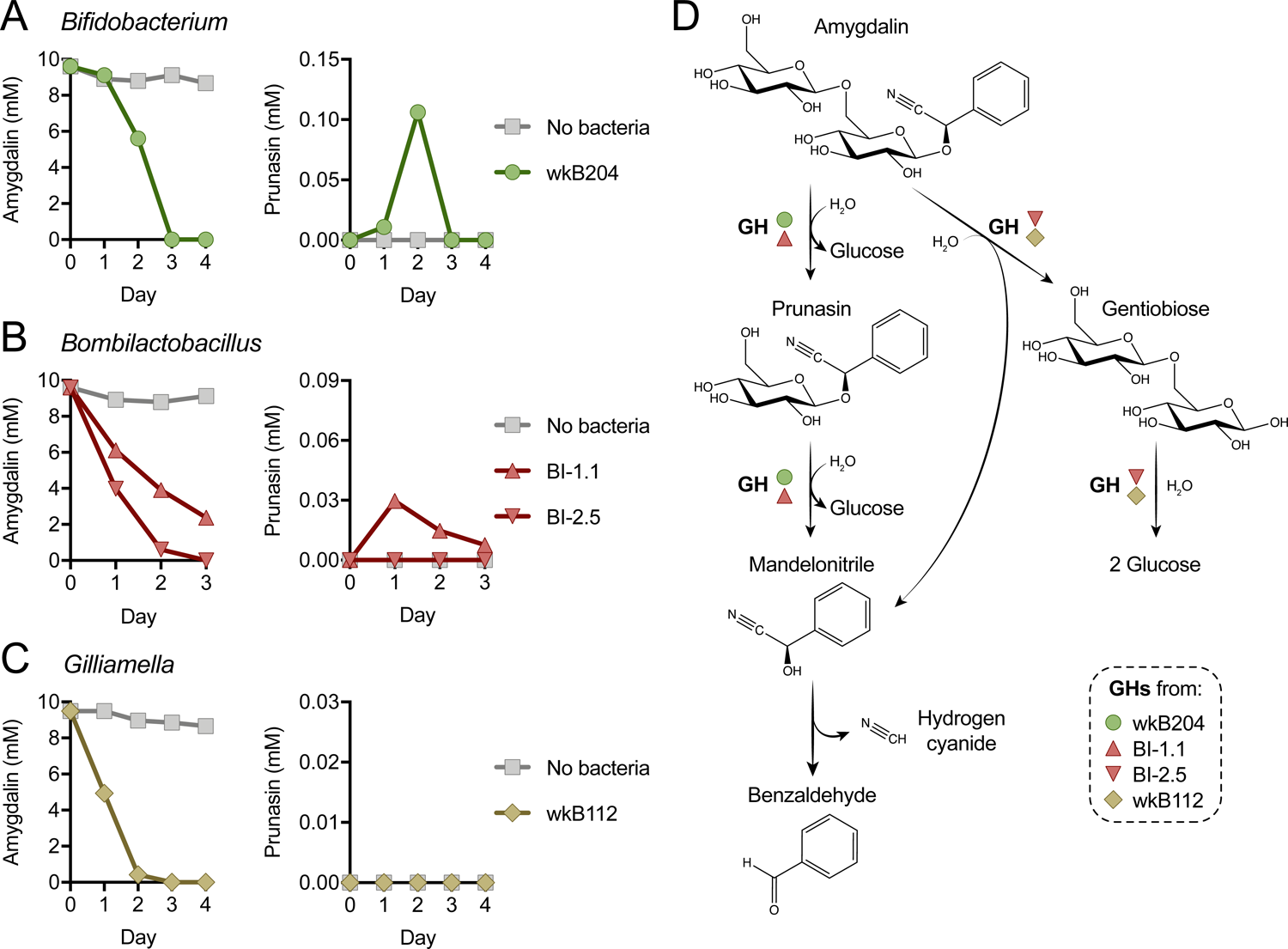
Mechanism of amygdalin degradation by bee gut bacteria. Amygdalin and prunasin concentrations detected by LC-MS in spent-medium of 3- or 4-day old cultures of **(A)** *Bifidobacterium* strain wkB204, **(B)** *Bombilactobacillus* strains BI-1.1 and BI-2.5, and **(C)** *Gilliamella* strain wkB112. Concentrations were determined every day for 3 to 4 days. Controls consisted of medium with amygdalin but no bacteria. Only wkB204 and BI-1.1 produced prunasin as an intermediate. **(D)** Proposed mechanism of amygdalin degradation by different bacterial species in the bee gut.

### Characterizing an enzyme involved in amygdalin metabolism

After finding that specific strains from different bee gut bacterial species can degrade amygdalin, we focused on honey bee-associated *Bifidobacterium* strains to investigate the enzyme involved in this metabolism. First, we checked whether the enzyme is secreted or not. Large cultures of wkB204, wkB344 and wkB338 were grown for 5 days (Figure 3A), after which we performed biochemical assays with both spent medium and cell lysate of glucose- and amygdalin-grown cultures.

**Figure 3.**
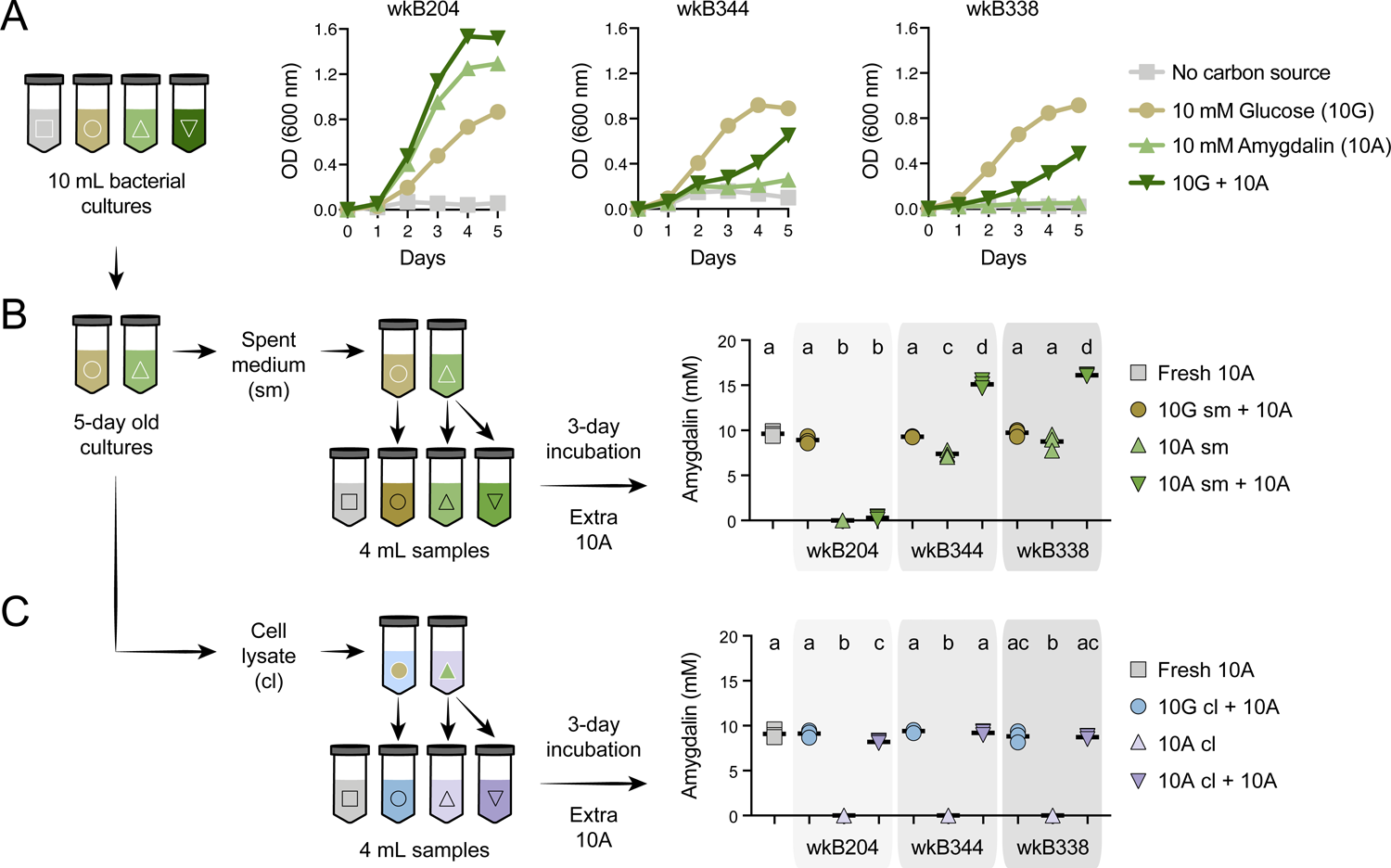
Amygdalin degradation in spent media and cell lysates of *Bifidobacterium* strains. **(A)** Bacterial growth curves of *Bifidobacterium* strains cultured in SDM without a carbon source, with 10 mM glucose (10G), with 10 mM amygdalin (10A), or with both 10 mM glucose and 10 mM amygdalin (10G + 10A) as carbon sources at 35 °C and 5% CO_2_. Experiments were performed in three biological replicates. Each data point represents the average optical density (600 nm) measured every day for 5 days. **(B-C)** For each strain, 10G and 10A grown cultures were separated into **(B)** spent medium (sm), originating from samples 10G sm and 10A sm, and **(C)** cell lysate (cl), originating from samples 10G cl and 10A cl. These samples were used to investigate amygdalin degradation by adding extra 10A to the samples. Controls consisted of 10A grown cultures without adding extra 10A and fresh SDM with 10A. Reactions were incubated at 35 °C and 5% CO_2_ for 3 days, after which amygdalin concentration was determined. Experiments were performed in three biological replicates. Groups with different letters are significantly different (*P* < 0.01, One-way ANOVA test followed by Tukey’s multiple-comparison test).

As observed in the previous experiment, wkB204 completely degraded amygdalin; we did not detect amygdalin in spent medium (10A sm, Figure 3B) or in cell lysate of amygdalin-grown cultures (10A cl, Figure 3C). To investigate whether the enzyme involved in amygdalin degradation was secreted, we added fresh amygdalin to sterile spent medium (10A sm + 10A) or to sterile cell lysate (10A cl + 10A) originating from amygdalin-grown cultures. After 3 days of incubation, we found full degradation of amygdalin in spent medium (10A sm + 10A) (Figure 3B), but only slight degradation in cell lysate (10A cl + 10A) (Figure 3C); this was compared to a control sample containing only medium and amygdalin (Fresh 10A). No amygdalin was detected in cell lysates of amygdalin-grown cultures, showing that amygdalin does not enter bacterial cells (10A cl, Figure 3C). Moreover, we detected prunasin in both spent medium and cell lysate of amygdalin-grown cultures supplemented with amygdalin (10A sm + 10A and 10A cl + 10A, respectively) (Figure S2).

For comparison, these assays were also performed for wkB344 and wkB338 cultures. Some amygdalin degradation was observed for the spent medium of wkB344 amygdalin-grown cultures, but this degradation was much less than that observed for wkB204 (Figure 3B-C). We also investigated enzyme production and activity in the absence of amygdalin, by adding amygdalin to spent medium and cell lysate from glucose-grown cultures (10G sm + 10A and 10G cl + 10A, Figure 3B-C). None of the strains was able to significantly degrade amygdalin under these conditions.

Since spent medium of wkB204 amygdalin-grown cultures achieved full degradation of amygdalin, we decided to characterize the secreted enzyme involved in the degradation. Spent media from wkB204 cultures, grown with either amygdalin or glucose, were processed to obtain concentrated protein extracts (Figure 4A). Protein profiles were first obtained by SDS-PAGE gel and showed that amygdalin-grown cultures had a distinct secretome when compared to glucose-grown cultures (Figure 4B). Then, samples were submitted to proteomics analysis, which confirmed the expression differences, as we found 107 proteins secreted in higher abundance in amygdalin-grown cultures and 131 proteins secreted in higher abundance in glucose-grown cultures (*P* < 0.05, t-test followed by Benjamini-Hochberg procedure to control for false discovery, Figure 4C). Several significantly upregulated proteins in amygdalin-grown cultures are associated with carbohydrate metabolism (Supplementary Dataset 1). Interestingly, we detected a highly expressed enzyme belonging to the glycoside hydrolase family 3 (WP_254476944) only in amygdalin-grown cultures (Figure 4C), suggesting its involvement in the observed degradation. Other studies have demonstrated that specific bacterial or fungal GH3 enzymes can degrade amygdalin (37–40).

**Figure 4.**
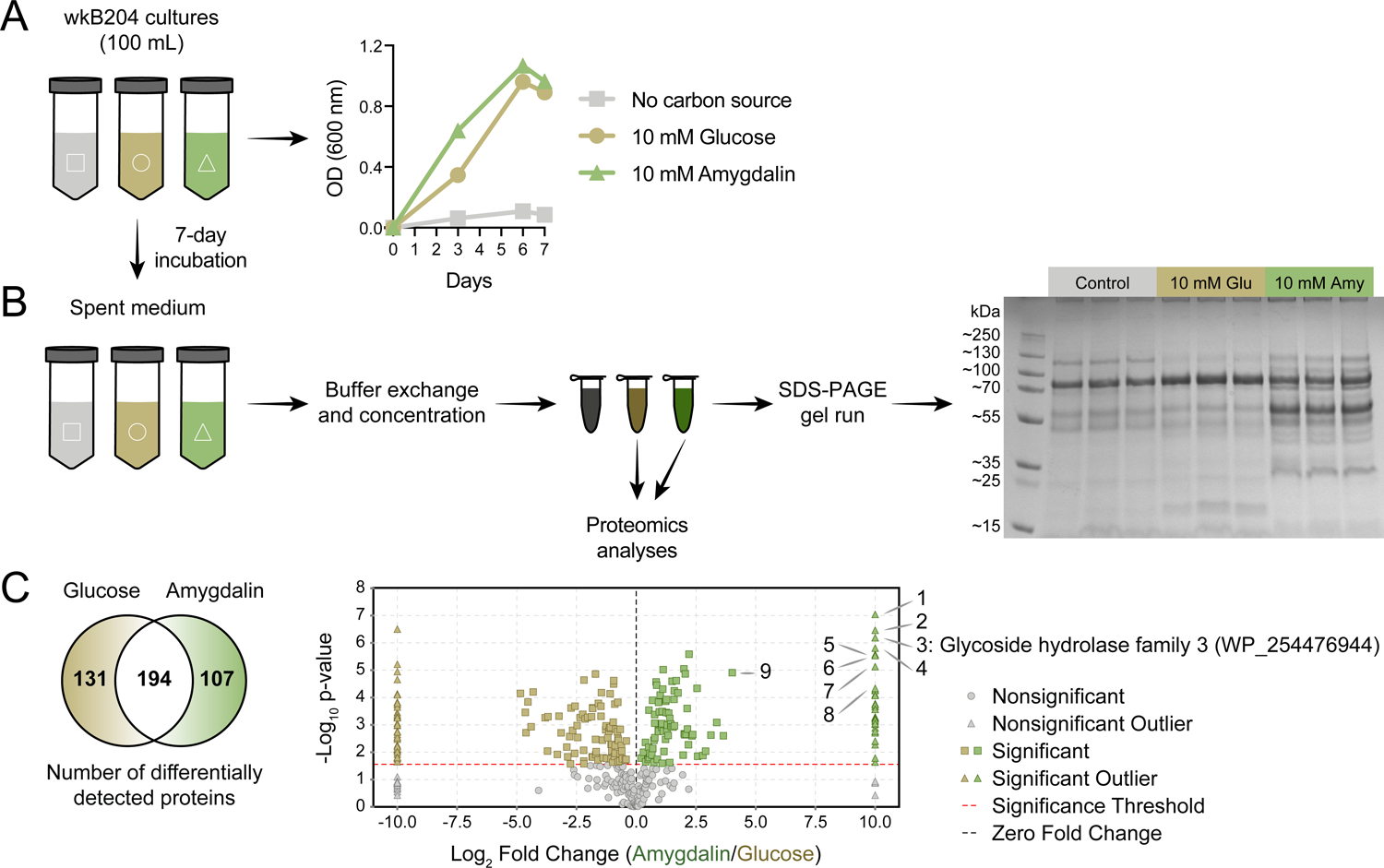
Identification of an amygdalin degrading enzyme from *Bifidobacterium*. (A) Large-scale culture of *Bifidobacterium* strain wkB204 in SDM without a carbon source, with 10 mM glucose, or with 10 mM amygdalin at 35 °C and 5% CO_2_. Experiments were performed in three biological replicates and each data point represents the average optical density (600 nm) measured every day for 7 days. **(B)** Spent medium concentration for running on a SDS-PAGE gel. **(C)** Venn diagram and volcano plot showing the number of differentially expressed proteins in spent medium of glucose- or amygdalin-grown cultures. Numbers in the volcano plot: 1: Alpha/beta fold hydrolase (WP_254477374), 2: Nucleoside hydrolase (WP_254477231), 3: Glycoside hydrolase family 3 (WP_254476944), 4: Beta-galactosidase (WP_254477161), 5: Alpha-mannosidase (WP_254477012), 6: Nudix hydrolase (WP_254477413), 7: MFS transporter (WP_254476943), 8: Alpha-L-fucosidase (WP_254477430), 9: Glycoside hydrolase family 30 (WP_254477160). (*P* < 0.05, T-test followed by Benjamini-Hochberg procedure to control for false discovery rate). Figure 4 **– source data 1.** SDS-PAGE gel run for cultures of *Bifidobacterium* strain wkB204. From left to right, columns represent: (1) PageRuler™ Plus Prestained Protein Ladder; (2–10) Supernatants of cultures (1–4) grown in the absence of a carbon source, (5–7) in the presence of 10 mM glucose as sole carbon source, or (8–10) in the presence of 10 mM amygdalin as sole carbon source. Each sample (30 μL) was mixed with 5 μL of 6X SDS gel-loading buffer (0.35 M Tris-Cl pH 6.8, 10% w/v SDS, 0.012% w/v bromophenol blue, 30% v/v glycerol, 0.6 mM dithiothreitol), denatured at 100 °C for 5 minutes, then run on a Bolt^TM^ 4-12% Bis-Tris Plus, 1.0 mm, protein gel at 200 V for 22 minutes.

### GH3 gene expression in *Bifidobacterium* strains

We used this wkB204 GH3 (WP_254476944) as a query to search a customized database of proteins from bee gut bacteria, including 22 bee-associated *Bifidobacterium* strains. Ten other *Bifidobacterium* strains encode a GH3 in their genomes with a high sequence similarity to the wkB204 GH3 (Figure S3). Intriguingly, these included a GH3 from wkB344 (WP_121913979), which did not grow in the presence of amygdalin *in vitro*.

To determine why this GH3 does not enable wkB344 to use amygdalin as a carbon source, we investigated whether this enzyme is expressed in cultures, and used wkB204 and wkB338 cultures as controls for presence and absence of GH3 activity, respectively (Figure 5A). In the presence of glucose as the sole carbon source, strains wkB204 and wkB344, but not wkB338, express the GH3 gene (Figure 5B). When cultivated in the presence of amygdalin as the sole carbon source, only wkB204 shows elevated expression of GH3 transcripts (Figure 5B), which correlates with the ability of this strain to degrade amygdalin *in vitro*. No elevation in expression was evident for wkB344 (Figure 5B), and the levels of GH3 produced by wkB344 in glucose-grown cultures did not result in observable amygdalin degradation when incubated in 10 mM amygdalin (Figure 3C).

**Figure 5.**
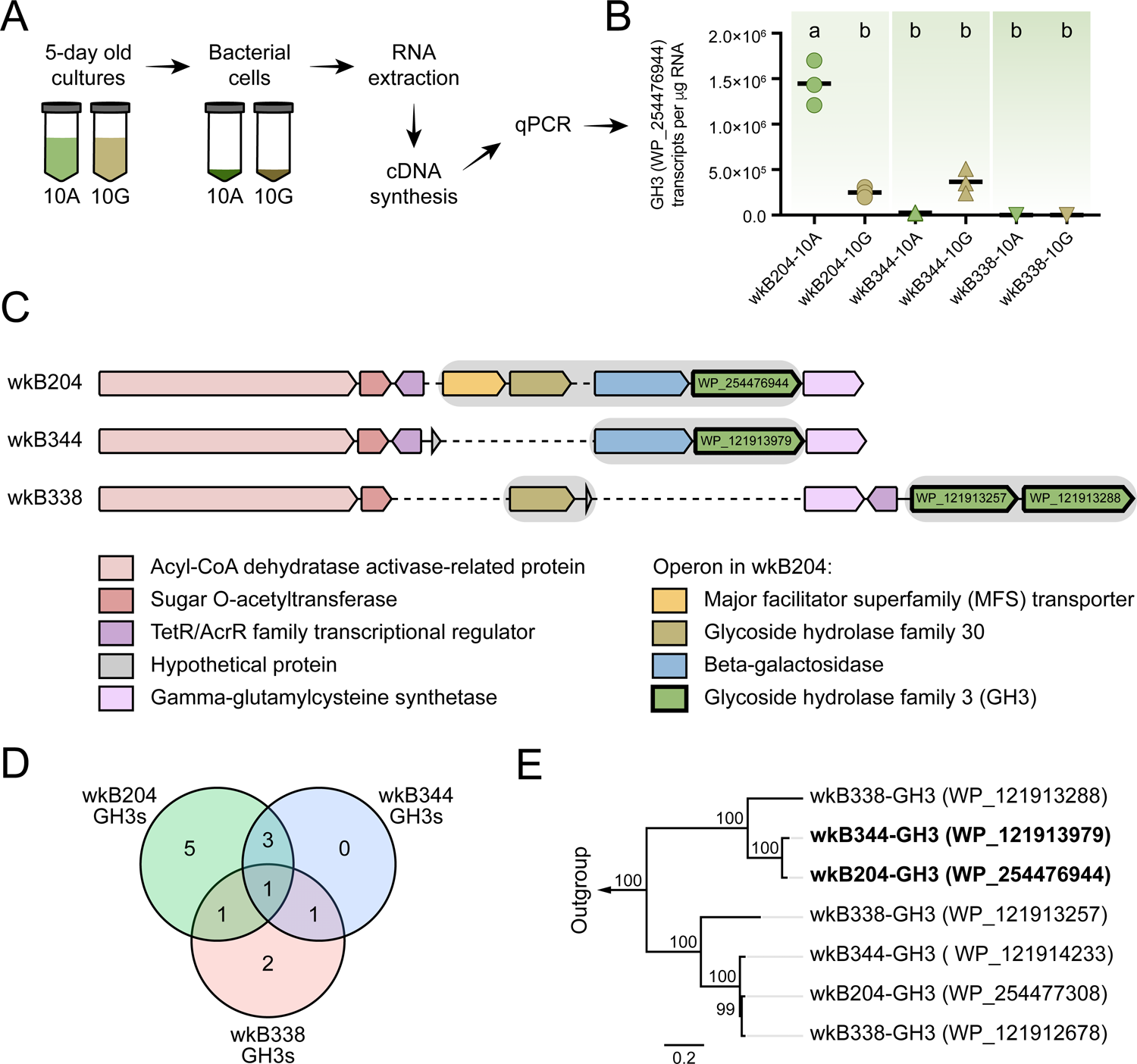
GH3 gene expression in *Bifidobacterium* cultures. **(A)** RNA extraction and cDNA synthesis from cultures of *Bifidobacterium* strains wkB204, wkB344 and wkB338. **(B)** qPCR data for the transcript levels of GH3 in cells of *Bifidobacterium* strains cultured in the presence of 10 mM glucose (10G) or 10 mM amygdalin (10A). Experiments were performed in three biological replicates. Groups with different letters are significantly different (*P* < 0.01, One-way ANOVA test followed by Tukey’s multiple-comparison test). **(C)** The genomic region containing the GH3 gene with high sequence similarity in wkB204 and wkB344. The corresponding region is included for wkB338 for comparison. Gray shading indicates operons. Dashed lines indicate regions not present in the genome. **(D)** Venn diagram showing the number of GH3s shared between the strains with amino acid similarity to other annotated GH3s according to the NCBI inference database. **(E)** Phylogenetic analysis for the GH3s found in the genomic regions shown in Figure 5C. Outgroup is represented by two amygdalin-degrading GH3s isolated from *Rhizomucor miehei* strain RmBglu3B (AIY32164.1) and *Talaromyces cellulolyticus* strain Bgl3B (GAM39187.1).

The wkB204 GH3 (WP_254476944), which is overexpressed in amygdalin-grown cultures, is encoded in an operon containing four other genes: a major facilitator superfamily transporter (WP_254476943), a glycoside hydrolase family 30 (WP_254477160), and a beta-galactosidase (WP_254477161) (Figure 5C). These were also overexpressed in the presence of amygdalin, based on proteomics data (Figure 5C). The wkB344 GH3 (WP_121913979) is also encoded in an operon, but a beta-galactosidase (WP_121914045) is the only other gene in the operon (Figure 5C), as predicted by the operon-mapper webserver (41).

According to the dbCAN meta server for automated CAZyme annotation, the genomes of these three *Bifidobacterium* strains encode multiple GH3s: wkB204 encodes 10 distinct GH3s, while wkB344 and wkB338 encode 5 distinct GH3s each (Supplementary Dataset 2) (42, 43). Based on the NCBI inference database and amino acid similarity to other annotated GH3s, these three strains have some GH3s highly similar in amino acid sequence and probably similar in function (Figure 5D and Table S1), as noted for wkB204-GH3 (WP_254476944) and wkB344-GH3 (WP_121913979) (Figure 5E).

### *Bifidobacterium* strains also degrade prunasin

To investigate whether *Bifidobacterium* strains can also degrade prunasin, we performed an additional *in vitro* experiment in which *Bifidobacterium* strains wkB204, wkB344 and wkB338 were grown in 10 mM glucose in SDM in the presence of 0.1 mM prunasin (Figure 6A). Under these conditions, all strains grew in the presence of prunasin (Figure 6B) and degraded it (Figure 6C). For comparison, we also checked growth in the presence of 0.1 mM amygdalin (Figure 6D) and found that not only wkB204, but also wkB344 degraded amygdalin (Figure 6E). The lack of growth and, consequently, degradation observed before for this strain is probably due to the high concentration of amygdalin provided in previous cultures (10 mM or 100 mM).

**Figure 6.**
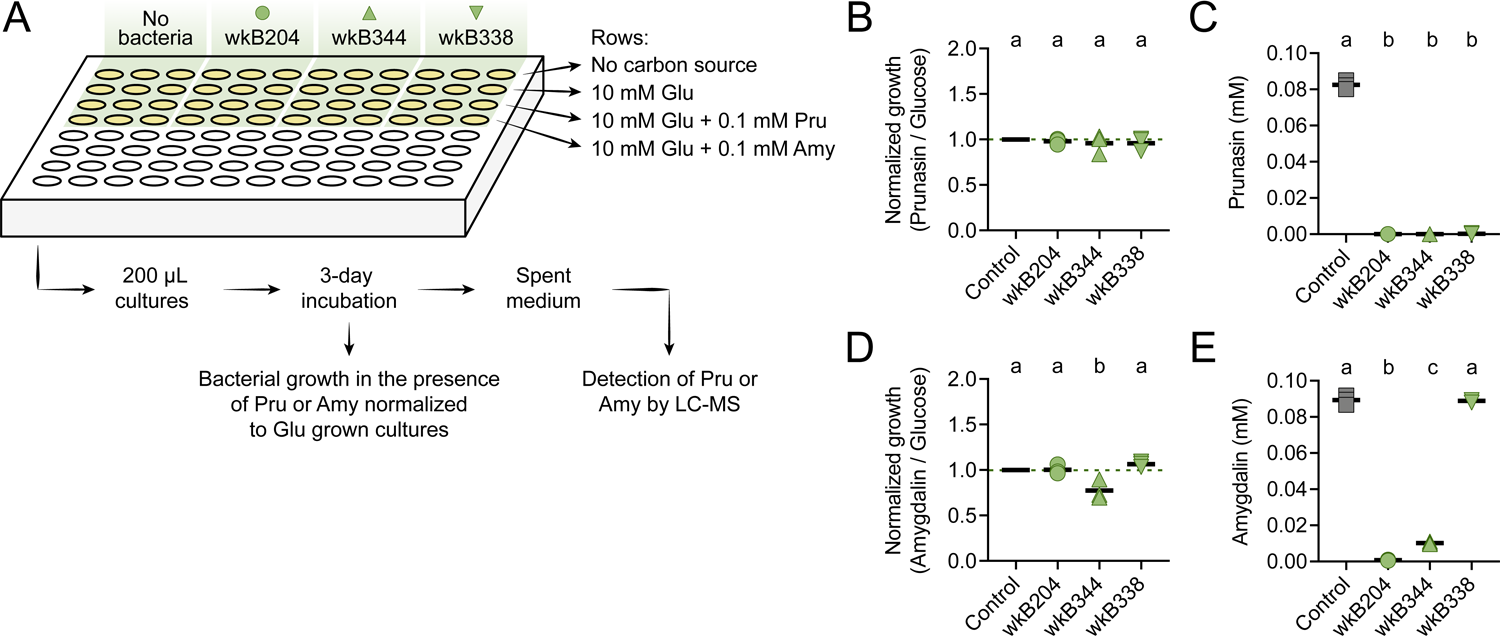
Prunasin degradation by bee gut-associated *Bifidobacterium* strains. **(A)** Experimental design. **(B)** Bacterial growth and **(C)** prunasin degradation after 3 days of incubation in the presence of 0.1 mM prunasin. **(D)** Bacterial growth and **(E)** amygdalin degradation after 3 days of incubation in the presence of 0.1 mM amygdalin. Experiments were performed in three biological replicates. Groups with different letters are significantly different (*P* < 0.01, One-way ANOVA test followed by Tukey’s multiple-comparison test).

### *Escherichia coli* expressing the GH3 enzyme produces prunasin

To confirm the ability of the *Bifidobacterium* GH3 enzyme to degrade amygdalin and/or prunasin, we cloned and expressed the GH3 gene from *Bifidobacterium* strains wkB204 (WP_254476944) or wkB344 (WP_121913979) in *E. coli* (Figure 7A-B). Cell lysates of transformed *E. coli* expressing GH3 were incubated in the presence of 0.1 mM amygdalin or 0.1 mM prunasin (Figure 7C). After 5 days of incubation, we observed amygdalin degradation (Figure 7D) followed by prunasin production (Figure 7E) for *E. coli* cell lysates expressing either wkB204-GH3 or wkB344-GH3, but not for *E. coli* transformed with an empty plasmid, suggesting that both enzymes can degrade amygdalin into prunasin. When the cell lysates were incubated in the presence of prunasin, only a small amount of prunasin was degraded (Figure 7F), suggesting that this enzyme, under the tested conditions, still can degrade prunasin, but to a lesser extent. These findings show that this *Bifidobacterium*-related GH3 enzyme can degrade amygdalin into prunasin, and potentially prunasin into mandelonitrile, and may be responsible for the degradation patterns observed for *Bifidobacterium* strain wkB204 when cultured in the presence of amygdalin.

**Figure 7.**
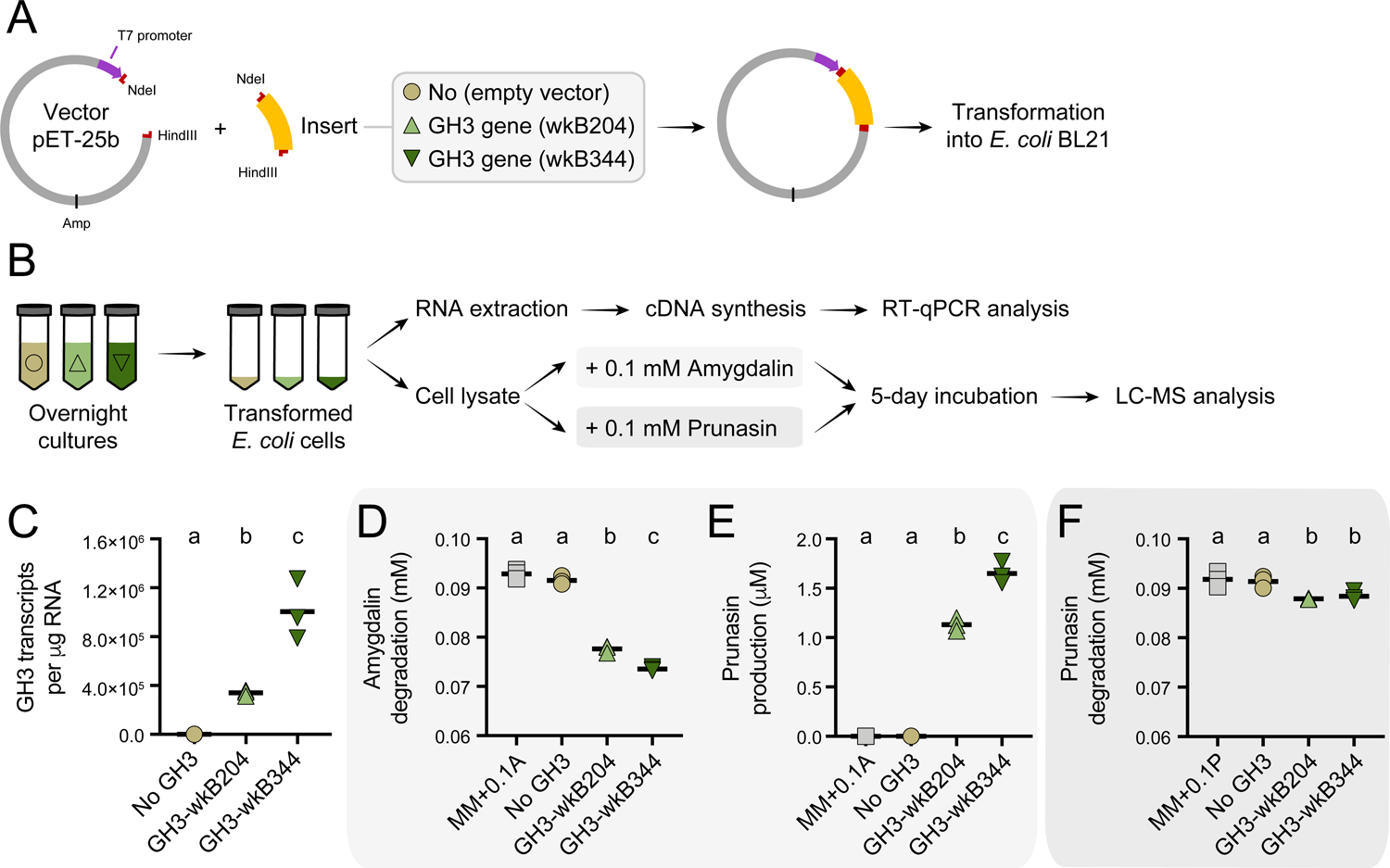
Heterologous expression of *Bifidobacterium* GH3 enzyme in *Escherichia coli*. **(A)** *E. coli* Rosetta BL21 competent cells were transformed with the vector pET-25b carrying the gene that encodes the wkB204-GH3 or wkB344-GH3, or only the empty vector as a control. **(B)** Bacterial cells from overnight cultures were lysed to extract RNA and investigate the expression levels of cloned genes by RT-qPCR. In parallel, bacterial cells from similar overnight cultures were lysed and used in incubation assays with 0.1 mM amygdalin or 0.1 mM prunasin in minimal medium at 37 °C. Samples were submitted for LC-MS analysis along with amygdalin and prunasin standards. **(C)** Transcript levels of *Bifidobacterium*-related GH3 genes expressed in *E. coli*. **(D)** Amygdalin degradation and **(E)** prunasin production levels after 5 days of incubation in the presence of 0.1 mM amygdalin. **(F)** Prunasin degradation levels after 5 days of incubation in the presence of 0.1 mM prunasin. Experiments were performed in three biological replicates. Groups with different letters are significantly different (*P* < 0.01, One-way ANOVA test followed by Tukey’s multiple-comparison test).

### Host and symbionts contribute to amygdalin degradation

We also investigated amygdalin degradation *in vivo*. To that end, we performed experiments with bees lacking a microbiota (microbiota-deprived or MD), colonized with a conventional microbiota (CV), or monocolonized with *Bifidobacterium* strains wkB204 or wkB344 (Figure 8A). Bees were hand-fed 5 μL of 1 mM amygdalin in sucrose syrup, or only sucrose syrup. Amygdalin was detected in different compartments of the bee body, including the midgut (M) and the body carcass without the gut (B) of MD, CV, and monocolonized bees (Figure 8B). In the hindgut (H) samples, amygdalin was detected for MD and wkB344-monocolonized bees but not for CV and wkB204-monocolonized bees (Figure 8B). Total amygdalin concentration was significantly lower in CV bees and wkB204-monocolonized bees when compared to control bees not treated with amygdalin, but spiked with 5 μL of 1 mM amygdalin during the extraction protocol (Figure 8C). Interestingly, prunasin was only detected in the midgut and hindgut of MD bees (Figure 8D-E). These findings demonstrate the role of the microbiota in amygdalin degradation, as amygdalin concentration is reduced in CV bees and prunasin does not accumulate in the guts of CV or monocolonized bees. These findings also provide evidence that bees themselves can degrade amygdalin, but that this degradation is partial, since prunasin accumulates in the guts of MD bees. Therefore, the presence of the microbiota contributes to continued amygdalin and prunasin degradation in the bee gut. It remains unknown whether this complete degradation, which potentially leads to the production of toxic hydrogen cyanide, is beneficial or detrimental to the bees.

**Figure 8.**
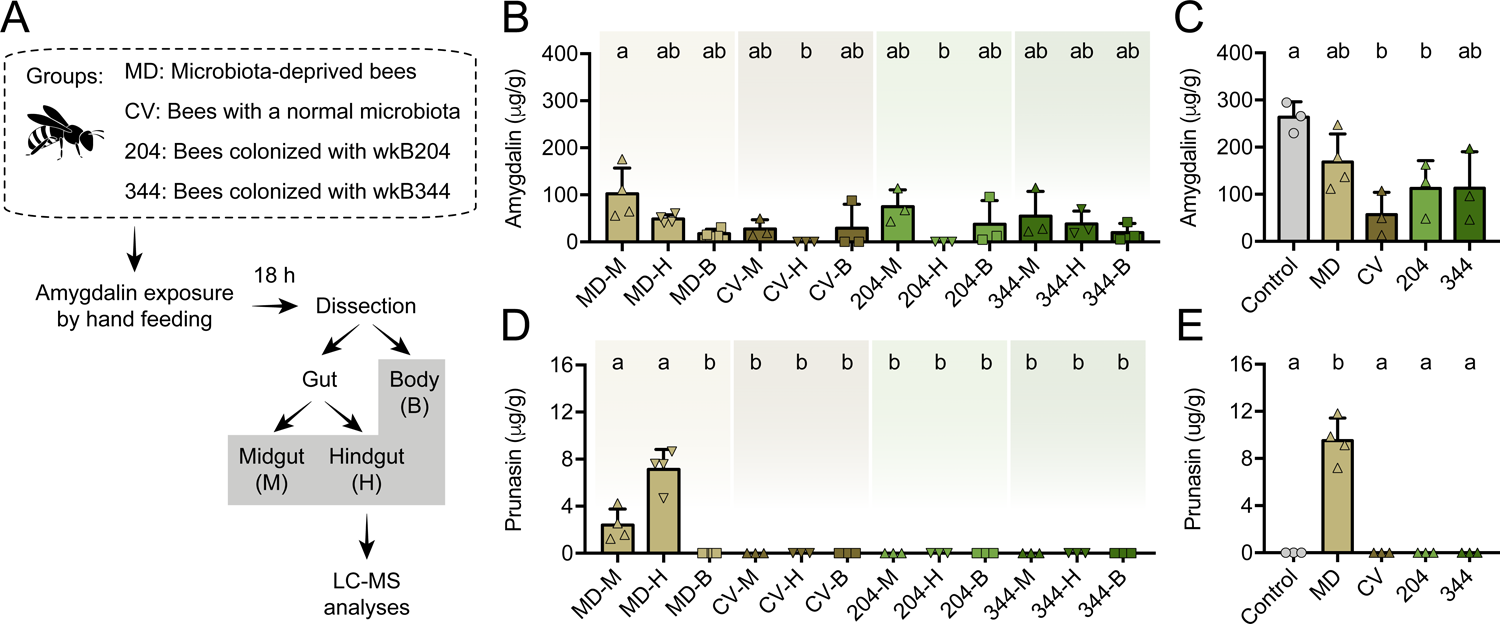
Amygdalin metabolism in honey bees. **(A)** Five-day old bees either lacking a microbiota (microbiota deprived, MD, n = 4), with a normal microbiota (conventionalized, CV, n = 3), or monocolonized with *Bifidobacterium* strains wkB204 (n = 3) or wkB344 (n = 3), were exposed to 5 μL of 1 mM amygdalin and after 24 hours dissected to determine the concentrations of **(B)** amygdalin and **(D)** prunasin in different bee body compartments (midgut: M, hindgut: H, and body without gut: B) by LC-MS. **(C)** Amygdalin and **(E)** prunasin concentrations detected in M, H and B samples were summed for each group and compared to a control group of unexposed bees that were mixed with 5 μL of 1 mM amygdalin at the beginning of sample processing. Groups with different letters are significantly different (*P* < 0.05, One-way ANOVA test followed by Tukey’s multiple-comparison test).

### Honey bees tolerate environmental concentrations of amygdalin

Honey bees exposed to environmental concentrations of amygdalin, ranging from 0.01 mM to 1 mM, did not exhibit increased mortality or dysbiosis (Figure S4A-B). We did not find any significant changes in gut microbial composition (Figure S4C-D) or abundance (Figure S4E) of amygdalin-treated bees when compared to untreated bees, which is consistent with other studies (29).

## Discussion

Pollinators such as bees are constantly exposed to nectar metabolites, which may exert varied effects on their health. In this study, we performed *in vitro* experiments with bee gut bacterial symbionts to investigate their ability to metabolize the plant toxin amygdalin, as well as *in vivo* experiments with bees to assess the contribution of the host and the microbiota to amygdalin degradation. Once consumed by bees, amygdalin can be broken down in the bee gut, but whether this degradation is via host or pollen-derived glycoside hydrolases (21, 23) or through activity of the microbiota, has been unclear. We found that members of the bee gut microbiota can degrade amygdalin and its intermediate prunasin *in vitro*. Specific strains of *Bifidobacterium*, *Bombilactobacillus* and *Gilliamella* isolated from *Apis mellifera*, *Bombus impatiens*, and *Apis cerana*, respectively, were able to degrade amygdalin *in vitro*. In some cases, the pathway led to the production of the intermediate prunasin; in others it did not.

We identified a GH3 in *Bifidobacterium* strains that contributes to amygdalin and prunasin metabolism *in vitro*. There are other studies showing that bacterial or fungal derived GH3s can degrade amygdalin. For example, the Gram-positive bacterium *Cellulomonas fimi* encodes a GH3 with activity against β-1,6-linked glycosides (37), similar to the linkage found in the structure of amygdalin (Figure 2D). Degradation of amygdalin by GH3s isolated from *Rhizomucor miehei* (38) and *Talaromyce leycettanus* (40), or by related extracellular enzymes from *Aspergillus niger*, have also been observed (39). Moreover, different species of mammalian gut-associated *Bifidobacterium* strains can grow in the presence of amygdalin, potentially due to the production of GH1 or GH3 enzymes (44). In our case, heterologous expression of wkB204-GH3 (WP_254476944) or wkB344-GH3 (WP_121913979) into *E. coli* also led to amygdalin degradation, but to a lesser extent than what was observed for the original host. To accurately quantify the lower degradation rates in transformed *E. coli*, we used 0.1 mM amygdalin solutions (Figure 7). Potentially, there are other carbohydrate digestive enzymes encoded by *Bifidobacterium* that contribute to amygdalin or prunasin metabolism (Figure 4C), or the *Bifidobacterium* host perform specific posttranslational modifications on this enzyme that are not achieved during heterologous expression in *E. coli* (45). Moreover, it seems this GH3 is secreted by *Bifidobacterium* strain wkB204, but, for cloning purposes, we expressed it in *E. coli* without a signal sequence for secretion, and, therefore, performed assays with cell lysates, which may not be optimal for characterizing enzyme activity. These features may have masked the potential activity of this GH3, if this is the main enzyme involved in amygdalin degradation by *Bifidobacterium* strains. Moreover, other hydrolases were identified in the proteomics data, such as apha/beta fold hydrolase (WP_254477374), nucleoside hydrolase (WP_254477231), beta-galactosidase (WP_254477161), alpha-mannosidase (WP_254477012), nudix hydrolase (WP_254477413), alpha-L-fucosidase (WP_254477430), and glycoside hydrolase family 30 (WP_254477160) (Figure 4C). Although not reported in the literature, some of these could potentially be involved in amygdalin degradation It is important to note that colonization by the normal microbiota decreases the pH to about 5 in the bee gut (46), and low pH can favor the degradation of amygdalin into prunasin, then prunasin into mandelonitrile, which can undergo spontaneous degradation at acidic pH to give benzaldehyde and hydrogen cyanide (47). Therefore, the presence of the microbiota itself, without the action of glycoside hydrolases, could favor the degradation of amygdalin into prunasin and potentially lead to the production of hydrogen cyanide.

As shown in other studies, nectar and pollen metabolites can be chemically transformed when passing through the gut, and this may influence their effects on the host and/or the microbiota (48, 49). To identify the potential contribution of both the host and the microbiota to amygdalin degradation, we performed *in vivo* experiments in which microbiota-deprived bees and microbiota-colonized bees were exposed to amygdalin for a short period of time. We found that microbiota-deprived bees can degrade amygdalin, but only partially, leading to accumulation of prunasin in gut compartments. In contrast, prunasin accumulation was not observed in microbiota-colonized bees or in bees monocolonized with specific *Bifidobacterium* strains, and amygdalin degradation was higher in these groups when compared to microbiota-deprived bees. This suggests that members of the microbiota, besides contributing to amygdalin degradation, can also efficiently degrade prunasin and potentially release the final products of amygdalin metabolism, such as hydrogen cyanide. This could potentially increase the side effects of amygdalin byproducts on bees or on the microbiota, since hydrogen cyanide can be toxic to aerobic organisms (9, 18). However, similar to other studies, we did not detect increased mortality rates or changes in microbial community abundance and composition for bees exposed to amygdalin (17, 29). Interestingly, we also found amygdalin in bee carcasses with the gut removed, suggesting that amygdalin is absorbed systemically by bee cells. This has been observed in other studies in which amygdalin was found in the bee hemolymph after oral ingestion (18, 49).

Studies in other animals have also investigated the roles of the microbiota on amygdalin degradation. Studies in rats, for example, have found higher concentrations of hydrogen cyanide in the blood of microbiota-colonized rats than in microbiota-deprived rats (10) or antibiotic treated rats (11) after oral ingestion of amygdalin. However, intravenous administration of amygdalin seems not to lead to hydrogen cyanide formation (47), which could be correlated to the lack of toxicity observed in honey bees after amygdalin injection into the hemolymph (18). These studies suggest that the gut microbiota is a major factor driving amygdalin degradation and hydrogen cyanide release in the guts of animals.

Although bees and their native microbiota seem to tolerate relatively high doses of amygdalin, parasites that commonly inhabit the bee gut may not fare as well. There is increasing evidence that metabolites in nectar and pollen, even those considered toxic in some cases, can improve pollinator health at specific concentrations by controlling or reducing parasite loads (16, 29, 50–52). Indeed, bees tend to forage on specific plants as a means of reducing colony pathogen loads (51). For example, honey bees from hives treated with amygdalin exhibited decreased levels of infection by the parasite *Lotmaria passim* and some viruses (29). In contrast, this does not seem to be the case for *Crithidia* infection in bumble bees, whose loads are not reduced after amygdalin exposure (53), as the parasite is not susceptible to amygdalin (54). Other plant metabolites, such as the essential oil thymol, can reduce *Nosema* spore loads in honey bees (50, 52). Therefore, the extent of protection may depend on the exposure level and on the parasite being exposed.

More recently, a few studies have brought attention to activities of plant-derived compounds within the host gut, which can be modulated by the host and/or the microbiota, to lead to the final metabolite activity (48). The recently described plant metabolite callunene, from heather nectar, can play a prophylactic role in preventing *Crithidia* infections in bumble bees, especially when parasite cells are present in the crop. However, callunene cannot play a therapeutic role in infected hosts, because host metabolism inactivates it before it reaches the hindgut where *Crithidia* usually establishes (48). In our study, on the other hand, we show that both the host and the microbiota contribute to the metabolism of amygdalin. Future studies could investigate whether hydrogen cyanide released into the gut may bring any positive effects to bees infected by specific parasites. It is known that the microbiota plays an important role in protection against infections (55) and that *Nosema* infection, for example, can lead to dysbiosis (56–58), but further studies are required to fully comprehend the mechanisms through which gut microbial communities deal with plant metabolites and the consequences for host fitness.

Our study shows the relevance of the microbiota for the metabolism of plant toxins, using amygdalin as an example of a toxin that is cooperatively metabolized by the host and the microbiota. Whether this full metabolism is beneficial, neutral, or harmful to bees is not yet known. A few other experiments have also investigated the roles of the microbiota in the metabolism of other plant metabolites. Kešnerová et al. (36) demonstrated that honey bees monocolonized with strains of *Bifidobacterium*, *Bombilactobacillus* or *Lactobacillus* can metabolize flavonoid glycosides. Indeed, genomic analyses have shown that the genomes of strains from these bacterial groups contain a diverse set of carbohydrate processing genes, including glycosidase hydrolases that could be involved in the cleavage of sugar residues (27, 28). We found strain variation for amygdalin metabolism, and such variation likely influences processing of other dietary secondary metabolites. Strain level variations in the microbiome could, consequently, affect parasite persistence or establishment. Perturbation of the honey bee gut microbiota by pesticides and/or heavy metals (59–61) could indirectly affect parasite success in the bee gut through changes in the metabolism of secondary metabolites by the microbiome.

## Methods

### Chemicals, media, and solutions

Amygdalin was obtained from Chem-Impex International, Inc. (catalog number: 22029, lot number: 002681-16112001). Prunasin was obtained from Toronto Research Chemicals, Inc. (catalog number: P839000, lot number: 6-EQJ-155-1). A semi-defined medium (SDM, for recipe see Table S2 and (62)) was used to culture *Bifidobacterium* and *Bombilactobacillus* strains. The nutrient-rich medium Insectagro DS2 (Corning, Inc., catalog number: 13-402-CV, lot number: 12818007) was used to culture *Gilliamella* strains. The Difco Lactobacilli MRS broth (BD, Inc., catalog number: 288130, lot number: 9211338) was used to culture *Lactobacillus* strains. Luria-Bertani (LB) or a minimal medium (MM, for recipe see Table S3 and (63)) was used to culture transformed *E. coli* strains. For experiments with bacterial isolates, a 1 M amygdalin solution was prepared by dissolving 4.57 g amygdalin in 10 mL of culture medium, then diluted to final concentrations of 0.1, 1, 10 or 100 mM in the same culture medium. Also, a 5 mM prunasin solution was prepared by dissolving 5 mg prunasin in 3,387 μL sterile water, then an aliquot was transferred to SDM or MM to a final concentration of 0.1 mM prunasin. For experiments with honey bees, a 10 mM amygdalin solution was prepared by dissolving 45.74 mg amygdalin in 10 mL sterile water, then diluted to final concentrations of 0.01 mM, 0.1 mM or 1 mM with filter-sterilized 0.5 M sucrose syrup and provided to bees in cup cages.

### Isolation and characterization of *Bifidobacterium* strains

*Bifidobacterium* strains wkB204, wkB344, and wkB338 were isolated from fresh guts of *Apis mellifera* workers from hives kept at UT-Austin (August 2014). Guts were homogenized in 10% PBS and cultured on Heart Infusion agar at 35 °C and 5% CO_2_ for 3–5 days. Genomic DNA was extracted from overnight cultures, as in (64). The wkB204 genome was sequenced on the Illumina MiSeq platform from 2×150-bp paired-end libraries at the SeqCenter (Pittsburgh, PA) and assembled using CLC Genomics Workbench 5.5 (Qiagen). The wkB344 and wkB338 genomes were previously reported in (31).

### Isolation and characterization of *Bombilactobacillus* and *Lactobacillus* strains

Bee gut-associated bacterial strains were isolated from fresh guts of commercial (strains BI-2.5, BI-1.1) and wild-caught (strain BI-4G) *Bombus impatiens* workers, preserved guts of *Bombus appositus* (strain LV-8.1) and *Bombus occidentalis* (strain OCC3) workers. Wild *B. impatiens* were collected in New Haven, CT, USA (August 2013); *B. appositus* and *B. occidentalis* were collected in Logan, UT, USA (July 2013); commercial *B. impatiens* were obtained from BioBest (Romulus, MI, USA). Guts and feces were homogenized in 10% PBS and cultured in MRS broth at 35 °C and 5% CO_2_ for 3–5 days, then plated on MRS agar and incubated at 35 °C and 5% CO_2_. Several passages on MRS agar were required to achieve pure isolates. Genomic DNA was extracted from overnight cultures, as in (64).

The BI-2.5 genome was sequenced and closed using Pacific Biosciences technology at the Yale Center for Genome Analysis. Indel errors were corrected with Pilon using Illumina MiSeq reads – 150-bp single-read libraries sequenced at the GSAF, UT-Austin (65). The other four genomes were sequenced on the Illumina MiSeq platform from 2×300-bp paired-end libraries at the GSAF, UT-Austin and assembled using CLC Genomics Workbench 5.5 (QIAGEN). All genomes were annotated with the Rapid Annotation using Subsystem Technology (RAST) server (66). Strains BI-2.5, BI-1.1 and LV-8.1 are most related to *Bombilactobacillus bombi* (67, 68), while strains OCC3 and BI-4G are most related to *Lactobacillus bombicola* (69, 70).

*Bombilactobacillus mellifer* strain Bin4N (DSM 26254) was obtained from the Leibniz Institute, Germany and is a honey bee isolate (67, 71).

*Lactobacillus* strains HB-1, HB-2, HB-C2, and HB-D10 were isolated from fresh guts of *Apis mellifera* workers from hives kept at UT-Austin (August 2017). Guts were homogenized in 10% PBS and cultured in MRS broth at 35 °C and 5% CO_2_ for 3–5 days. Aliquots of bacterial cultures were plated on MRS agar and incubated at 35 °C and 5% CO_2_. Several passages on MRS agar were required to achieve pure isolates. Sequencing the 16S rRNA gene showed that these isolates corresponded to the bee-restricted cluster that contains *Lactobacillus melliventris*.

*Lactobacillus* strains wkB8 and wkB10 were previously isolated from the guts of *Apis mellifera* (72).

### Isolation and characterization of *Gilliamella* strains

*Gilliamella* strains were previously isolated from the guts of *Apis dorsata* (wkB112, wkB178, wkB108), *Apis cerana* (wkB308) or *Apis mellifera* (M6-3G, M1-2G, wkB7, wkB1) (31).

### Exposure of bee gut bacteria to amygdalin

All strains were initially cultured in Heart Infusion Agar (Criterion, USA, catalog number: C5822, lot number: 491030) with 5% Defibrinated Sheep Blood (HemoStat Laboratories, USA, lot number: 663895-2) at 35 °C and 5% CO_2_ for 3–5 days, then colonies were transferred to proper liquid media to obtain enough bacterial mass for *in vitro* experiments.

Strains of *Bifidobacterium* (wkB204, wkB338, wkB344) and *Bombilactobacillus* (BI-1.1, BI-2.5, LV-8.1, Bin4N) were cultured in SDM (Table S2) at 35 °C and 5% CO_2_ overnight. Optical density (OD) of each bacterial culture was measured at 600 nm, and the cells were diluted to an OD of 0.5 in SDM. Ten-microliter aliquots of each bacterial suspension were transferred in three biological replicates to 96-well plates containing 190 μL SDM with no carbon sources, 10- or 100-mM amygdalin, 10- or 100-mM glucose, or 10- or 100-mM amygdalin and glucose as carbon sources. Controls consisted of three biological replicates of 200 μL SDM with similar carbon sources, but without bacterial suspension. The plates were incubated in a plate reader (Tecan) at 35 °C and 5% CO_2_, and OD was measured at 600 nm after 72 hours.

Strains of *Gilliamella* (wkB112, wkB178, wkB108, wkB308, M6-3G, M1-2G, wkB7, wkB1) were cultured in a nutrient-rich medium, Insectagro DS2 (Corning Inc.), at 35 °C and 5% CO_2_ overnight. OD of each bacterial culture was measured at 600 nm, and the cells were diluted to an OD of 0.5 in Insectagro. Ten-microliter aliquots of each bacterial suspension were transferred in three biological replicates to 96-well plates containing 190 μL Insectagro with 10- or 100-mM amygdalin, or without amygdalin. Controls consisted of three biological replicates of 200 μL Insectagro with similar carbon sources, but without bacterial suspension. The plates were incubated in a plate reader (Tecan^®^) at 35 °C and 5% CO_2_ and OD was measured at 600 nm every 3 hours for 72 hours.

Strains of *Lactobacillus* nr. *melliventris* (HB-1, HB-2, HB-C2, HB-D10, wkB8, wkB10, BI-4G, OCC3) were cultured in MRS broth at 35 °C and 5% CO_2_ overnight. OD of each bacterial culture was measured at 600 nm, and the cells were diluted to an OD of 0.5 in MRS. Ten-microliter aliquots of each bacterial suspension were transferred in three biological replicates to 96-well plates containing 190 μL MRS with 10 mM amygdalin, or without amygdalin. Controls consisted of three biological replicates of 200 μL MRS with similar carbon sources, but without bacterial suspension. The plates were incubated in a plate reader (Tecan^®^) at 35 °C and 5% CO_2_ and OD was measured at 600 nm every 3 hours for 72 hours.

At the end of the experiment, plates were centrifuged at 7000 rpm for 5 minutes, and spent medium was removed and filter-sterilized with a 0.22 μm filter. Samples were transferred to 1.5 mL microtubes and dried under vacuum using an Eppendorf Vacufuge (Eppendorf, USA). Later, they were resuspended in 1 mL LC-MS grade water and 100-fold diluted to be submitted for LC-MS analysis.

### Amygdalin degradation in spent media and cell lysates

*Bifidobacterium* strains wkB204, wkB344 and wkB338 were chosen to investigate the mechanism of amygdalin degradation. They were cultured in 5 mL of MRS broth at 35 °C and 5% CO_2_ overnight. OD was measured for each bacterial culture at 600 nm and diluted to an OD of 0.5 with SDM. Cells were washed twice with SDM. One hundred-microliter aliquots of each bacterial suspension were transferred in three biological replicates to 15 mL culture tubes containing 10 mL of SDM with 10 mM amygdalin, 10 mM glucose, or both, or without a carbon source. Samples were incubated at 35 °C and 5% CO_2_ and OD was measured at 600 nm every day for 5 days. At the end of the experiment, amygdalin- and glucose-grown cultures from each strain were centrifuged for 10 min at full speed, and spent medium was separated from the cell pellet.

Spent media of amygdalin- and glucose-grown cultures were filter-sterilized with a 0.22 μm filter, and 2.7 mL aliquots of each sample were transferred in three biological replicates to 15 mL culture tubes containing 0.3 mL of 100 mM amygdalin in SDM to investigate degradation by potential enzymes released into the media.

Bacterial cells of amygdalin- and glucose-grown cultures were washed three times with 1 mL SDM, then the supernatant was removed by centrifugation at full speed for 5 minutes. Washed cells were lysed with 1 mL of a bacterial protein extraction reagent (B-PER) solution, which consisted of 10 μL of 1 M MgCl_2_, 20 μL of 0.5 M phenylmethylsulfonyl fluoride (in methanol) and 9970 μL B-PER (Thermo Scientific, catalog number: 78248, lot number: LJ148147A). After 15 minutes, samples were centrifuged, filter-sterilized with a 0.22 μm filter, and 0.3 mL aliquots of each sample were transferred in three biological replicates to 15 mL culture tubes containing 0.3 mL of 100 mM amygdalin in SDM and 2.4 mL SDM to investigate degradation. Although the cell densities of wkB338 and wkB344 cultures were lower than that of wkB204 (Figure 3A), we were still able to collect and concentrate cells through centrifugation for the subsequent experimental steps. Normalization between samples was based on the volume of growth medium, and not on cell mass.

Spent medium and cell lysate of amygdalin-grown cultures only, and 10 mM amygdalin in fresh SDM were used as controls.

All samples were incubated at 35 °C and 5% CO_2_ for 3 days, after which they were 500-fold diluted and submitted for LC-MS analysis.

### Quantification of amygdalin in bacterial cultures

Diluted samples were analyzed using an Agilent 6546 Q-TOF LC-MS with an Agilent Dual Jet Stream electrospray ionization (ESI) source in negative mode. Chromatographic separations were obtained under gradient conditions by injecting 1 μL onto an Agilent RRHD Eclipse Plus C18 column (50 x 2.1 mm, 1.8 micron particle size) with an Agilent Zorbax Eclipse Plus C18 narrow bore guard column (12.5 x 2.1 mm, 5 micron particle size) on an Agilent 1260 Infinity II liquid chromatography system. The mobile phase consisted of eluent A (water + 0.1% formic acid) and eluent B (methanol). The gradient was as follows: held at 5% B from 0 to 1 min, 5% B to 30% B from 1 to 1.5 min, 30% B to 37% B from 1.5 to 9 min, 37% B to 95% B from 9 to 9.1 min, held at 95% B from 9.1 to 12 min, 95% B to 5% B from 12 to 12.1 min, and held at 5% B from 12.1 to 15 min. The flow rate was 0.4 mL/min. The sample tray and column compartment were set to 7 °C and 30 °C, respectively. The ion source settings were capillary voltage, 3,500 V; nozzle voltage, 2,000 V; fragmentor voltage, 180 V; drying gas and sheath gas temperature, 350 °C; drying gas flow, 10 L/min; sheath gas flow, 11 L/min; nebulizer pressure, 60 lb/in^2^. Q-TOF data was processed using Agilent MassHunter Qualitative Analysis software. Amygdalin (C_20_H_27_NO_11_) and prunasin (C_14_H_17_NO_6_) were observed in the samples with this LC-MS method as [M-H]^-^ at 456.1511 and 294.0983 Da, as well as [M+CH_3_COO]^-^ at 502.1566 Da and 340.1038 Da, with a retention time of 2.73 and 3.05 minutes, respectively. Amygdalin quantification was performed by preparing analytical curves using the area under the amygdalin extracted ion chromatogram peak (20 ppm extraction window) of the following standard solutions prepared from a 1 mM amygdalin stock solution in water: 0.078125, 0.15625, 0.3125, 0.625, 1.25, 2.5, 5, 10, 20, 30, and 40 μM amygdalin. Two analytical curves were prepared; one with the 6 six lower concentrations to calculate amygdalin concentration in 0.1- or 10-mM amygdalin cultures; another curve with the 5 higher concentrations to calculate amygdalin concentration in 100 mM amygdalin cultures. The linear equations obtained from these analytical curves were used to calculate the concentration of amygdalin in the samples. The concentrations obtained from the linear equation were corrected for the dilution factor. Prunasin quantification was performed similarly, by preparing an analytical curve using the area under the prunasin extracted ion chromatogram peak of the following standard solutions prepared from a 1 mM prunasin stock solution in water: 0.01953125, 0.0390625, 0.078125, 0.15625, 0.3125, 0.625, 1.25, 2.5, 5, and 10 μM prunasin. The linear equation obtained from this analytical curve was used to calculate the concentration of prunasin in the samples. The concentrations obtained from the linear equation were corrected for the dilution factor.

*Bifidobacterium* strain wkB204 was cultured in 5 mL of MRS broth at 35 °C and 5% CO_2_ overnight. OD was measured at 600 nm and adjusted to 0.5 with SDM. Bacterial cells were washed two times with SDM and resuspended in SDM. 200 μL aliquots were transferred in three biological replicates to 250 mL culture flasks containing 100 mL of SDM with 10 mM amygdalin, 10 mM glucose, or without a carbon source. Samples were incubated at 35 °C and 5% CO_2_ and OD was measured at 600 nm every day for 7 days. At the end of the experiment, samples were centrifuged at 7800 rpm for 10 min. Spent media were separated from bacterial cells and concentrated to about 10 mL in under vacuum using an Eppendorf Vacufuge (Eppendorf, USA). Then, samples were dialyzed three times in 1 L of exchange buffer (10% glycerol, 1mM MgCl_2_, 0.1 M NaCl, 1mM PMSF and 25 mM Tris pH 8), after which they were further concentrated with centrifugal concentrators (10 kDa MWCO, Millipore Sigma-Aldrich, USA) to a final volume of 1.5 mL. 30 μL of each concentrated sample were run on a Bolt 4-12% Bis-Tris Plus Gel (Thermo Scientific, catalog number: NW04120BOX, log number: 21022470). Then, concentrated samples from amygdalin- and glucose-grown cultures were submitted for proteomics analysis at the Proteomics facility, UT-Austin. The samples were digested with trypsin, desalted and run on the Dionex LC and Orbitrap Fusion 1 for LC-MS/MS with one hour run time and processed by the facility using PD 2.2 and Scaffold proteomics software (Proteome Software, Inc., Portland Oregon, version 5.1.2). For protein assignment, we used the amino acid sequences predicted from the wkB204 genome combined with a list of common contaminants for the searches. A basic Scaffold analysis was performed using a custom amino acid sequence database covering the genome of wkB204, a reference database for *Saccharomyces cerevisiae* (because of the yeast extract portion of the SDM used to grow this strain), as well as a list of common contaminants, using min protein: 0.1% false discovery rate (FDR).

### Blast search and phylogenetic analysis

A local blast was performed to search for homologous proteins of the glycoside hydrolase family 3 (GH3) that was detected in wkB204 amygdalin-grown cultures. We used the amino acid sequence of wkB204-GH3 as a query to search for homologous proteins in a custom database containing amino acid sequences of several published bee gut bacterial genomes, including 22 bee gut associated *Bifidobacterium* strains. We applied a query coverage high-scoring sequence pair percent of 90. wkB204-GH3 and homologous proteins were used to build a phylogenetic tree. Amino acid sequences were aligned using Muscle (73) and used to infer a maximum-likelihood phylogeny (LG model + Gamma4, 100 bootstrap replicates) with PhyML 3.1 (74) implemented in SeaView (75).

### GH3 gene expression in *Bifidobacterium* strains

100 μL of 0.5 OD cultures of *Bifidobacterium* strains wkB204, wkB344, and wkB338 were transferred in three biological replicates to SDM with 10 mM amygdalin or 10 mM glucose for a final volume of 10 mL. After 5 days, bacterial cultures were centrifuged to separate the supernatant from the cells. Total RNA was extracted from washed bacterial cells using the Quick-RNA Miniprep kit (Zymo Research, USA). To that end, bacterial cells were resuspended and lysed in 600 μL of RNA Lysis Buffer, and transferred to a capped vial containing 0.5 mL of 0.1-mm Zirconia beads (BioSpec Products, USA). Samples were bead-beaten for 2 × 30 s, centrifuged at 14,000 rpm for 30 s, and transferred to a new 1.5 mL microtube. After this step, extraction followed the protocol provided by Zymo Research. Final RNA samples were dissolved in 50 μL of water and stored at −80 °C. RNA concentrations were measured in a Qubit instrument and normalized to 200 ng/μL. Complementary DNA (cDNA) was synthesized using the qScript cDNA Synthesis Kit (QuantaBio, USA) following the manufacturer’s instructions, and stored at −20 °C. cDNA samples were 10-fold diluted to be used as templates for qPCR analyses.

Specific primers targeting a conserved 124 bp region in the GH3 gene found in *Bifidobacterium* strains wkB204 and wkB344 (B-GH3-F: 5’-CTACCGCAATCCCGACCT-3’, and B-GH3-R: 5’-CACCTCCTTGTCCACTCCC-3’) were designed and used to amplify total copies of GH3 gene transcripts in each sample on 384-well plates on a Thermo Fisher QuantStudio 5 instrument. Three technical replicates of 10 μL reactions were carried out for each sample with 5 μL iTaq Universal SYBR Green Supermix (Bio-Rad, USA), 0.05 μL (each) 100 μM primer, 3.9 μL H_2_O, and 1.0 μL template DNA. The cycling conditions consisted of an initial cycle of 50 °C for 2 min and 95 °C for 2 min, followed by 40 cycles of a two-step PCR of 95 °C for 15 s and 60 °C for 1 min. Quantification was based on standard curves from amplification of the cloned target sequence in the pGEM-T Easy vector (Promega, USA). Briefly, genomic DNA of *Bifidobacterium* strain wkB204 was used as a template to amplify the GH3 gene region of interest (124 bp) using the primers B-GH3-F and B-GH3-R. The purified amplicon was ligated into the pGEM-T Easy vector (Promega, USA). The recombined vector was purified and transformed into *E. coli* strain DH5-alpha competent cells via electroporation using the Gene Pulser Xcell Electroporation System (Bio-Rad, USA). The recombined vector was then isolated from an overnight culture using the Monarch Plasmid Miniprep Kit (New England BioLabs, USA), digested by the restriction enzyme ApaI (New England Biolabs, USA), purified, quantified in a Qubit 4 fluorometer (Invitrogen, USA) and the final concentration was adjusted so it could be used as a standard for qPCR reactions.

### Cloning and transformation experiments

*E. coli* strain DH5-alpha was used for gene cloning and *E. coli* strain Rosetta BL21 was used for heterologous expression. LB or MM (Table S3) supplemented with 100 μg/mL ampicillin were used for the cultivation. *E. coli* strains were always cultured at 37 °C overnight. The vector pET25b (Invitrogen, USA) was applied for cloning and expression. First, the vector pET25b-empty was transformed into *E. coli* DH5-alpha cells via electroporation using the Gene Pulser Xcell Electroporation System (Bio-Rad, USA). Positive transformants were screened on LB plates with 100 μg/mL ampicillin and by PCR amplification. An overnight culture was used to isolate the vector pET25b-empty (Monarch Plasmid Miniprep Kit, New England BioLabs, USA), which was then dephosphorylated with Antarctic Phosphatase (New England BioLabs, USA) to reduce recyclization. Genomic DNA of *Bifidobacterium* strains wkB204 and wkB344 were used as templates to amplify their respective amygdalin degrading GH3 enzymes by PCR. Specific primers, GH3-NdeI-F (5’-ttgtttaactttaagaaggagatatacatatggcatcaaggaagttgacagagg-3’) and GH3-HindIII-R (5’-agcccgtttgatctcgagtgcggccgcaagcttacccacggtcaccgtca-3’) were designed to amplify the whole gene encoding the amygdalin-degrading GH3 enzyme. The PCR products were purified and submitted for Sanger sequencing for confirmation. The purified vector pET25b-empty and the PCR product of wkB204-GH3 (or wkB344-GH3) were digested by the restriction enzymes NdeI and HindIII-HF (both from New England Biolabs, USA) and then ligated to construct the recombinant plasmid pET25b-wkB204-GH3 (or pET25b-wkB344-GH3). The sequence-verified recombinant plasmids were purified and transformed into *E. coli* Rosetta BL21 competent cells via electroporation using the Gene Pulser Xcell Electroporation System (Bio-Rad, USA). The empty plasmid was also transformed into *E. coli* Rosetta BL21 competent cells to be used as a control in the experiments. Positive transformants were screened on LB plates with 100 μg/mL ampicillin and by PCR amplification, and bacterial stocks were made from single cell, overnight cultures.

### GH3 gene expression in transformed *E. coli* strains

100 μL of 0.5 OD cultures of *E. coli* Rosetta BL21 cells carrying pET25b-empty, pET25b-wkB204-GH3 or pET25b-wkB204-GH3 were transferred in three biological replicates to 5 mL LB broth supplemented with 100 μg/mL ampicillin and 100 μg/mL isopropyl β-D-1-thiogalactopyranoside (IPTG). Bacterial cultures were grown overnight at 37°C, after which cells were separated from the supernatant by centrifugation. Total RNA was extracted from washed cells using the Quick-RNA Miniprep kit (Zymo Research, USA), cDNA was synthesized using the qScript cDNA Synthesis Kit (QuantaBio, USA), and qPCR was performed using the primers B-GH3-F and B-GH3-R and following the protocol described in the “GH3 gene expression in *Bifidobacterium* strains” section.

### Amygdalin and prunasin degradation in cell lysates of transformed *E. coli*

*In vitro* experiments were performed with transformed *E. coli* Rosetta BL21 cells carrying pET25b-empty, pET25b-wkB204-GH3 or pET25b-wkB204-GH3. To that end, transformants were grown overnight at 37 °C in LB supplemented with 100 μg/mL ampicillin and 100 μg/mL isopropyl β-D-1-thiogalactopyranoside (IPTG). OD was adjusted to 1 and cells were washed twice with MM (Table 2). 5 mL of 1 OD washed bacterial cultures were transferred to 5 mL Falcon tubes, centrifuged for 10 min at 7000 rpm, supernatant was removed, and cells were resuspended in 5 mL MM. Bacterial cells were centrifuged again and media was removed. Washed cells were lysed with 1 mL of B-PER solution, as described above, for 15 min at room temperature, after which 4 mL of MM was added.

Samples were filter-sterilized with a 0.22 μm filter and dialyzed in centrifugal concentrators (10 kDa MWCO, Millipore Sigma-Aldrich, USA) for 20 minutes. After dialysis, the final volume of concentrated samples was adjusted to 5 mL with MM. 0.5 mL aliquots of each sample were transferred in three biological replicates to 1.5 mL tubes containing 0.5 mL of 0.2 mM amygdalin or 0.2 mM prunasin in MM to investigate degradation. 0.1 mM amygdalin in fresh MM or 0.1 mM prunasin in fresh MM were used as controls. Samples were incubated at 37 °C for 5 days, after which they were 10-fold diluted and submitted for LC-MS analysis.

### *In vivo* experiment to investigate amygdalin degradation in the bee gut

Late-stage pupae (with eyes pigmented but lacking movement) of *Apis mellifera* female workers were aseptically removed from a brood frame from a hive kept at UT-Austin. Pupae were placed on Kimwipes in sterile plastic bins and placed in an incubator at 35 °C and ∼60% relative humidity to simulate hive conditions until emerging as adults. After three days, newly emerged workers (NEWs), which lack their normal microbiota, were transferred to cup cages containing sterile sucrose syrup and sterile bee bread. Approximately 400 NEWs were randomly divided into four groups which were fed sterile sucrose syrup and specific treatments as described below. Group 1 was exposed to sterile pollen, and therefore the bees remained as microbiota deprived. Group 2 was exposed to a fresh bee gut homogenate mixed with sterile pollen, and therefore the bees acquired the normal microbiota. The gut homogenate was prepared by aseptically pulling out the guts from 10 healthy workers from the same hive and mixing with equal proportions of 1x PBS and sterile sucrose syrup (5 mL total volume), and 200 μL of gut homogenate were transferred to sterile pollen and provided to the bees in each cup cage. Groups 3 and 4 were exposed to a *Bifidobacterium* wkB204 or wkB344 bacterial suspension, respectively. Each bacterial strain was cultured in SDM at 35 °C and 5% CO_2_ overnight. The 600 nm optical density (OD) of each bacterial culture was measured, cells were washed with 1x PBS, and diluted to a concentration of 0.5 OD in equal proportions of 1x PBS and sterile sucrose syrup. 200 μL of bacterial suspension were transferred to the bee bread provided to the bees in each cup cage. After 5 days, which is sufficient time for establishment of the gut microbiota (3), bees were transferred to 0.5 mL vials with tips cut off, then starved for 6 h, after which they were hand-fed with 5 μL of 1 mM amygdalin in sterile sugar syrup. They were kept in the same vial for 18 hours, after which they were frozen until further processing. The control group consisted of unexposed bees that were mixed with 5 μL of 1 mM amygdalin at the beginning of sample processing to determine the amount of amygdalin that would be detected before any degradation event could occur. Three bees from each group were thawed and aseptically dissected to obtain the following bee body compartments: midgut, hindgut, and bee body carcass. These samples were homogenized with 1 mL LC-MS grade water and submitted for LC-MS analyses.

### *In vivo* experiment to investigate the effects of amygdalin on the bee gut microbiota

A brood frame was collected from a honey bee hive at UT-Austin, transferred to a frame cage and placed in an incubator at 35 °C and ∼60% relative humidity to simulate hive conditions until adults emerged. 1-day old bees were randomly divided into four groups, each being treated with sucrose syrup, 0.01 mM amygdalin dissolved in sucrose syrup, 0.1 mM amygdalin dissolved in sucrose syrup, or 1 mM amygdalin dissolved in sucrose syrup. A gut homogenate (200 μL) was added to the bee bread provided to each cup cage, enabling colonization by the full gut microbiota, as in (59). 15 bees were sampled from each group after one week of treatment and stored at −80 °C. Each group consisted of 4 cup cages each containing 40 bees. Survival rates were monitored and dead bees were removed in a daily census.

### DNA extraction, qPCR analysis and 16S rRNA amplicon sequencing

Sampled honey bees were placed in sterile Falcon tubes and transferred to a freezer at −80 °C. DNA was extracted from individual guts, following a previously described protocol (64). Final DNA samples were 10-fold diluted to be used as templates for qPCR analyses, as described in (76), and for 16S rRNA library preparation and sequencing, as described in (77).

### Material availability and accession numbers

Bacterial strains are available by request from the Moran Lab. The complete genome sequence of strain BI-2.5 has been deposited at DDBJ/ENA/GenBank under the accession CP031513. The genome assemblies for strains BI-1.1, LV-8.1, BI-4G, L5-31, OCC3 and wkB204 have been deposited at DDBJ/ENA/GenBank under the accessions QOCR00000000, QOCS00000000, QOCU00000000, QOCT00000000, QOCV00000000 and JAFMNU020000000, respectively. 16S rRNA amplicon sequencing data are available at NCBI BioProject PRJNA865802.

## Acknowledgements

Thanks to the members of the Nancy Moran & Howard Ochman labs, especially Kim Hammond for maintaining hives, Eli Powell for lab assistance, and former member Margaret Steele for providing *Gilliamella* strains M1-2G and M6-3G. Thanks to Brianna Flynn for helping isolate *Lactobacillus* strains HB-1, HB-2, HB-C2, HB-D10, and Ryan Arnott for preparing 16S rRNA gene PCR amplicons from these strains for Sanger sequencing analyses. Thanks to Maria Person, Michelle Gadush and Peter Faull at the UT Austin Center for Biomedical Research Support Biological Mass Spectrometry Facility (RRID:SCR_021728) for processing protein samples and providing feedback on the raw data.

## Funding

This work was supported by the USDA National Institute of Food and Agriculture (grant number 2018-67013-27540).

## Supplementary material

### Supplementary Figures

**Figure S1.**
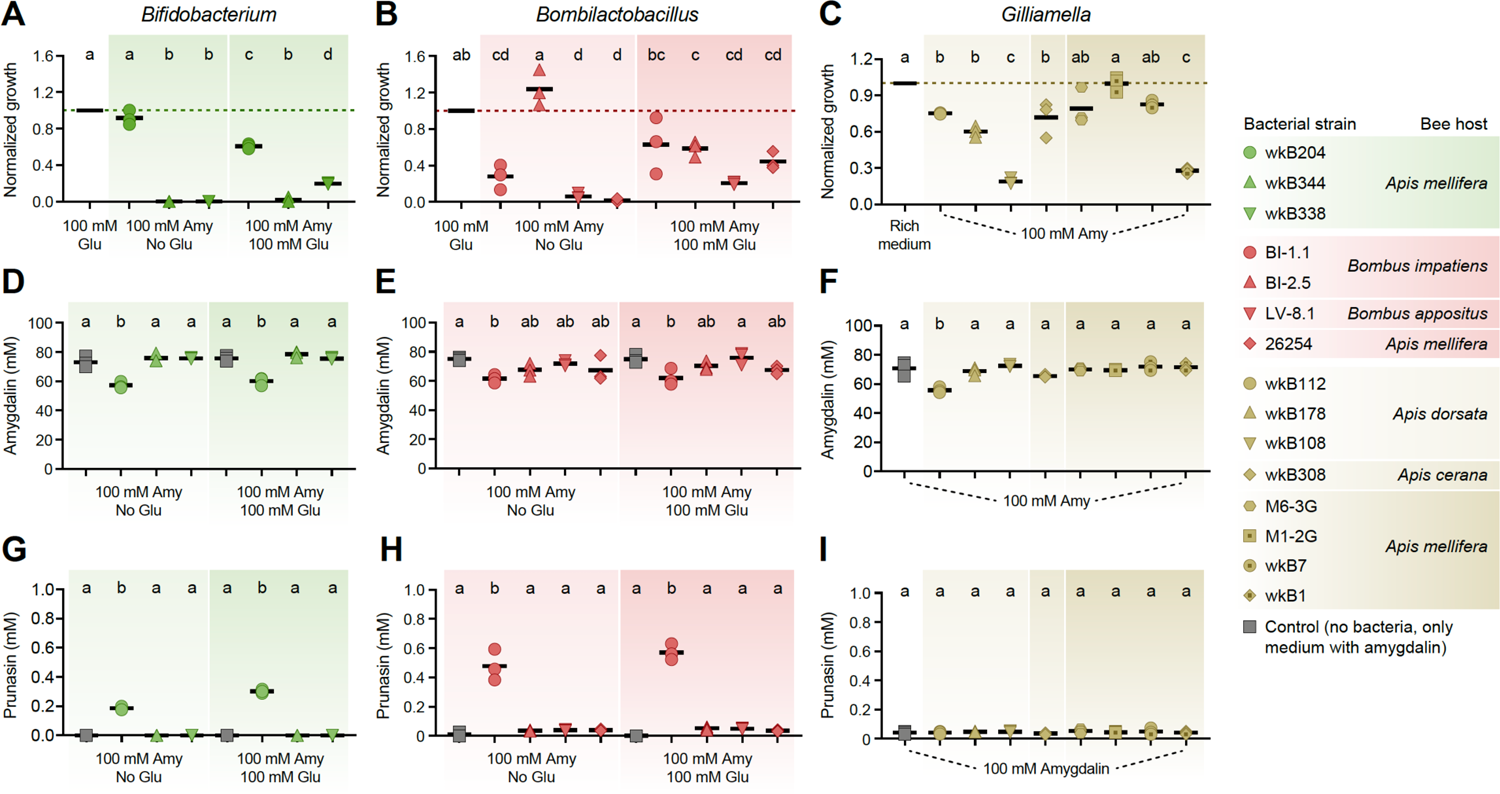
*In vitro* exposure of bee gut associated bacteria to 100 mM amygdalin. Growth of **(A)** *Bifidobacterium* and **(B)** *Bombilactobacillus* strains in semi-defined media in the presence of 100 mM amygdalin (or 100 mM amygdalin and 100 mM glucose) normalized to the bacterial growth in the presence of 100 mM glucose. Growth of **(C)** *Gilliamella* strains in nutritionally rich media in the presence of 100 mM amygdalin normalized to the bacterial growth in the absence of amygdalin. Bacterial growth was measured as optical density at 600 nm after 3 days of incubation at 35 °C and 5% CO_2_. **(D-F)** Amygdalin and **(G-H)** prunasin concentrations in spent medium of amygdalin (or amygdalin and glucose) grown cultures of *Bifidobacterium*, *Bombilactobacillus*, and *Gilliamella* strains, respectively. Controls consisted of media with amygdalin (or amygdalin and glucose) but no bacteria. Experiments were performed in three biological replicates. Groups with different letters are statistically significantly different (*P* < 0.01, One-way ANOVA test followed by Tukey’s multiple-comparison test).

**Figure S2.**
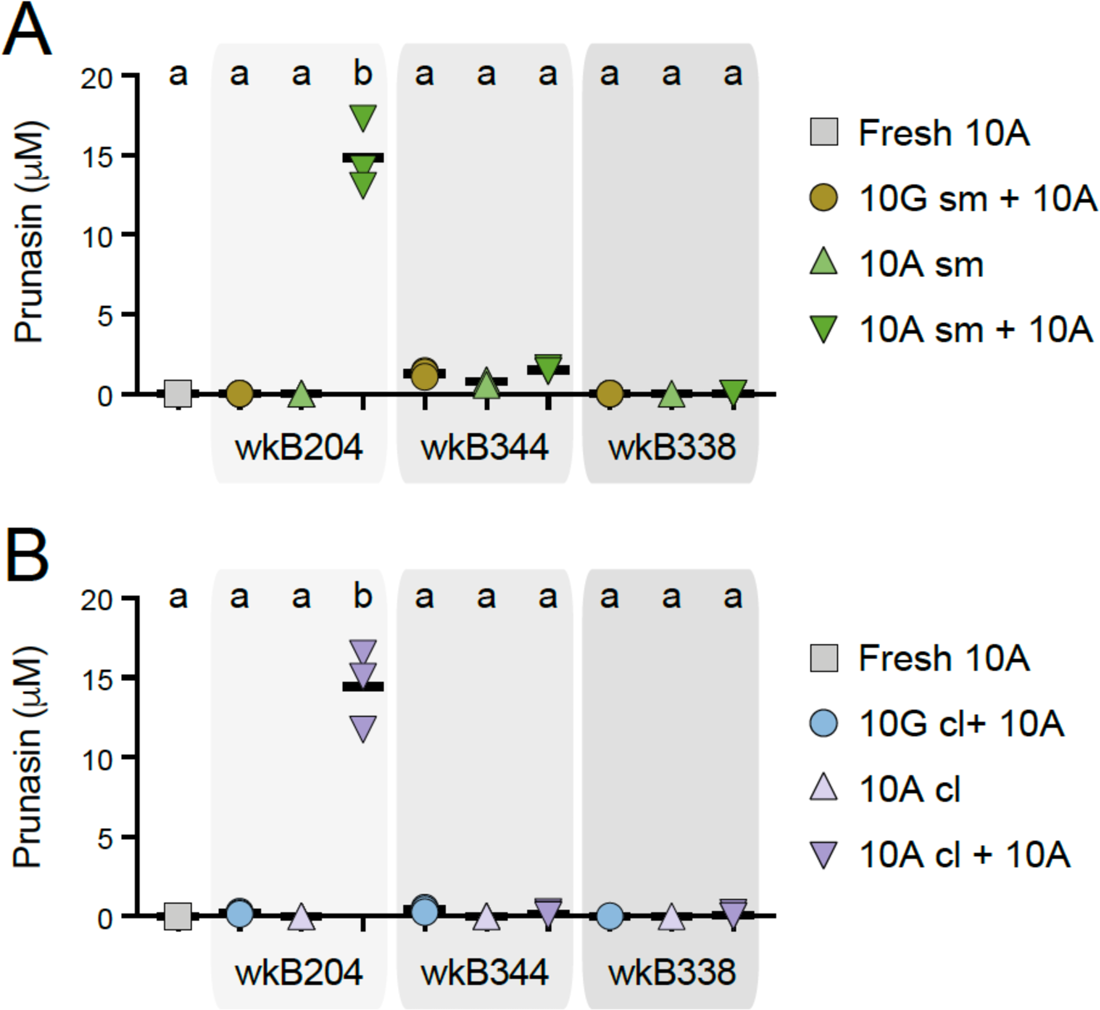
Prunasin concentrations in spent media and cell lysates of *Bifidobacterium* strains. *Bifidobacterium* strains were cultured in SDM without a carbon source, with 10 mM glucose (10G), with 10 mM amygdalin (10A), or with both 10 mM glucose and 10 mM amygdalin (10G + 10A) as carbon sources at 35°C and 5% CO_2_. For each strain, 10G and 10A grown cultures were separated into **(A)** spent medium (sm), originating the samples 10G-sm and 10A-sm, and **(B)** cell lysate (cl), originating the samples 10G-cl and 10A-cl. These samples were used to investigate prunasin release by adding extra 10A to the samples. Controls consisted of 10A grown cultures without adding extra 10A and fresh SDM with 10A. Reactions were incubated at 35 °C and 5% CO_2_ for 3 days, after which amygdalin concentration (see Figure 6) and prunasin release were determined. Experiments were performed in three biological replicates. Groups with different letters are statistically significantly different (P < 0.05, One-way ANOVA test followed by Tukey’s multiple-comparison test).

**Figure S3.**
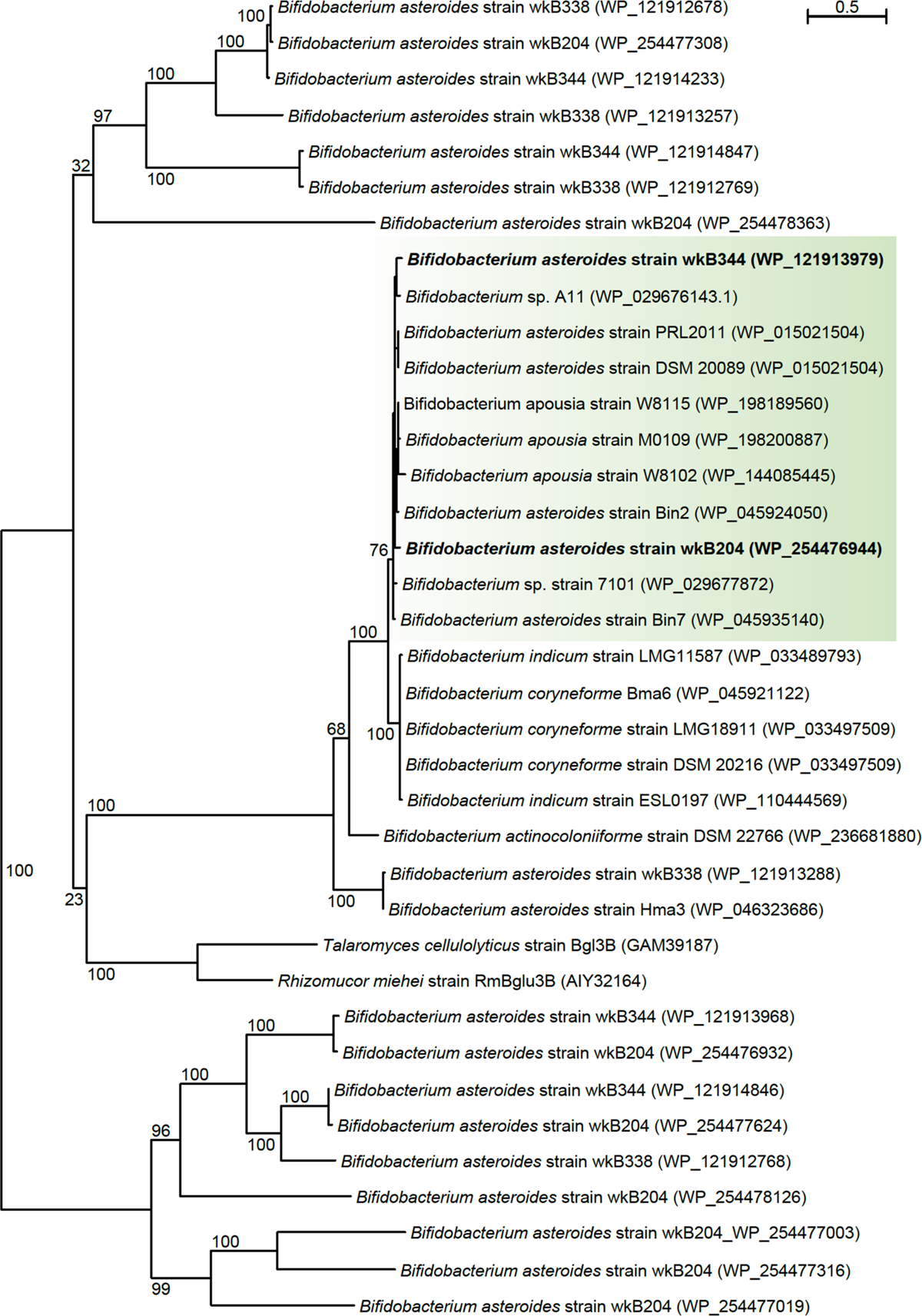
Maximum-likelihood phylogeny based on amino acid sequences of bee associated *Bifidobacterium* glycoside hydrolases with sequence homology to a glycoside hydrolase family 3 highly expressed in amygdalin grown cultures of *Bifidobacterium* strain wkB204 (PhyML 3.1, LG model + Gamma4, 100 bootstrap replicates).

**Figure S4.**
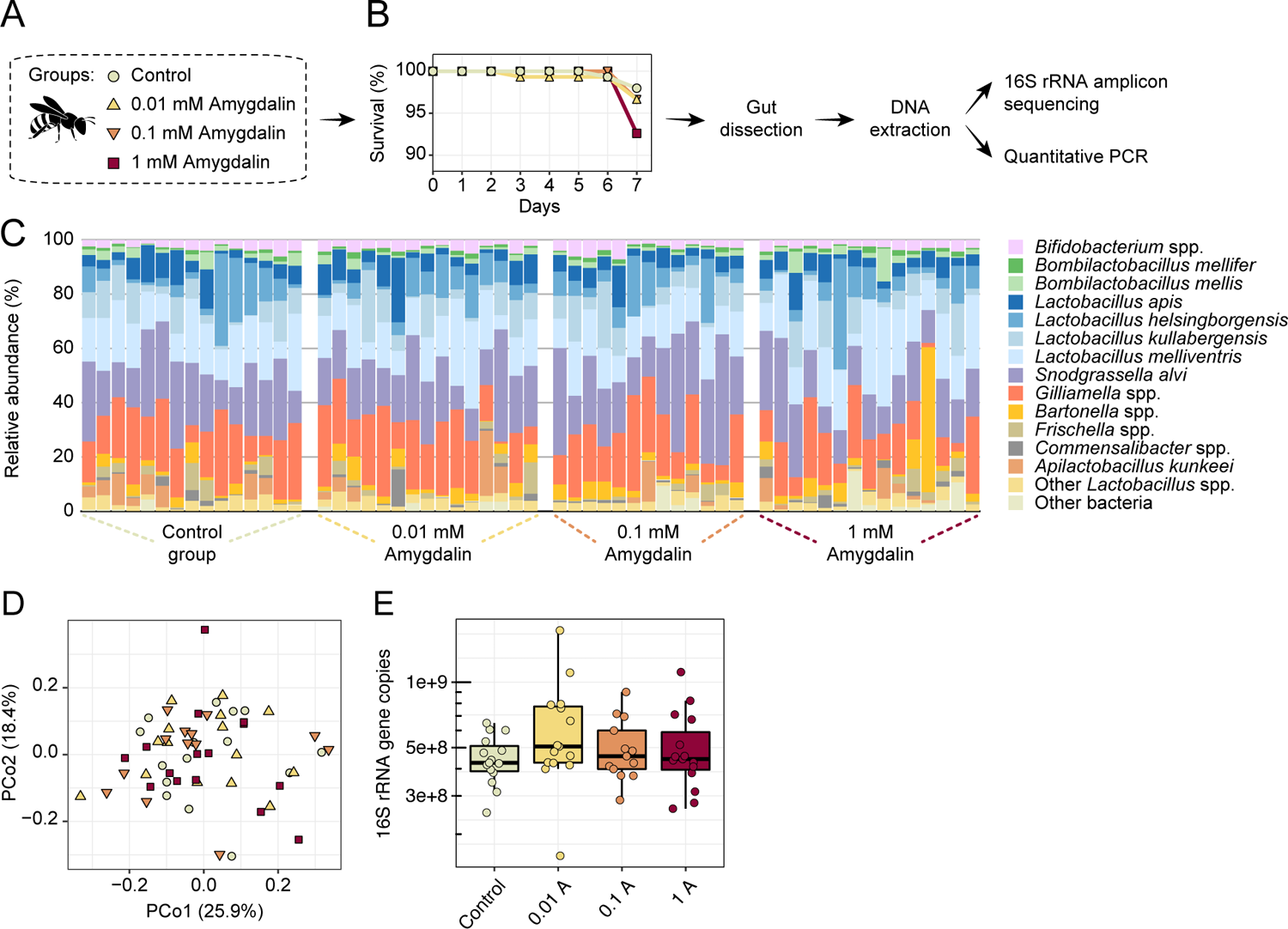
Amygdalin effects on the honey bee gut microbiota. (A) Experimental design and (B) survival rates of honey bees exposed to different concentrations of amygdalin. (C) Stacked column graphs showing the relative abundance of bee gut bacterial species in control bees (n = 15), 0.01 mM amygdalin (n = 15), 0.1 mM amygdalin (n = 13) and 1 mM amygdalin (n = 15) exposed bees. (D) Principal coordinate analysis of gut community compositions of control and amygdalin exposed bees using Bray-Curtis dissimilarity (Permanova test with 9,999 permuations; *P* > 0.5). (E) Boxplot of total bacterial 16S rRNA gene copies estimated by qPCR for control and amygdalin exposed bees. Box-and-whisker plots show high, low, and median values, with lower and upper edges of each box denoting first and third quartiles, respectively. No significant differences were observed in total bacterial abundance between control and amygdalin exposed bees (*P* > 0.05 Kruskal-Wallis test).

### Supplementary tables

**Table S1.** Glycoside hydrolases family 3 detected in the genomes of *Bifidobacterium* strains wkB204, wkB344 and wkB338. Protein ID refers to the unique identification of each GH3 in the NCBI Reference Sequence Database. Inference refers to the closest related GH3 present in the NCBI Reference Sequence Database. Similar colors indicate GH3s with similar amino acid sequence. This information was used to make the Venn Diagram in Figure 5D.

| Strain | GH3 loci number | Protein ID (NCBI RefSeq) | Inference (NCBI RefSeq) |
| --- | --- | --- | --- |
| wkB204 | 10 | WP_254476932 | WP_007147852 |
|  |  | WP_254476944 | WP_015021504 |
|  |  | WP_254477003 | WP_003842825 |
|  |  | WP_254477019 | - |
|  |  | WP_254477308 | WP_015022086 |
|  |  | WP_254477316 | - |
|  |  | WP_254477624 | WP_016461981 |
|  |  | WP_254477626 | WP_004221005 |
|  |  | WP_254478126 | - |
|  |  | WP_254478363 | - |
| wkB344 | 5 | WP_121913968 | WP_007147852 |
|  |  | WP_121913979 | WP_015021504 |
|  |  | WP_121914233 | WP_015022086 |
|  |  | WP_121914846 | WP_016461981 |
|  |  | WP_121914847 | WP_004221005 |
| wkB338 | 5 | WP_121912678 | WP_015022086 |
|  |  | WP_121912768 | WP_003838412 |
|  |  | WP_121912769 | WP_004221005 |
|  |  | WP_121913257 | WP_003839235 |
|  |  | WP_121913288 | WP_015450023 |

**Table S2.**
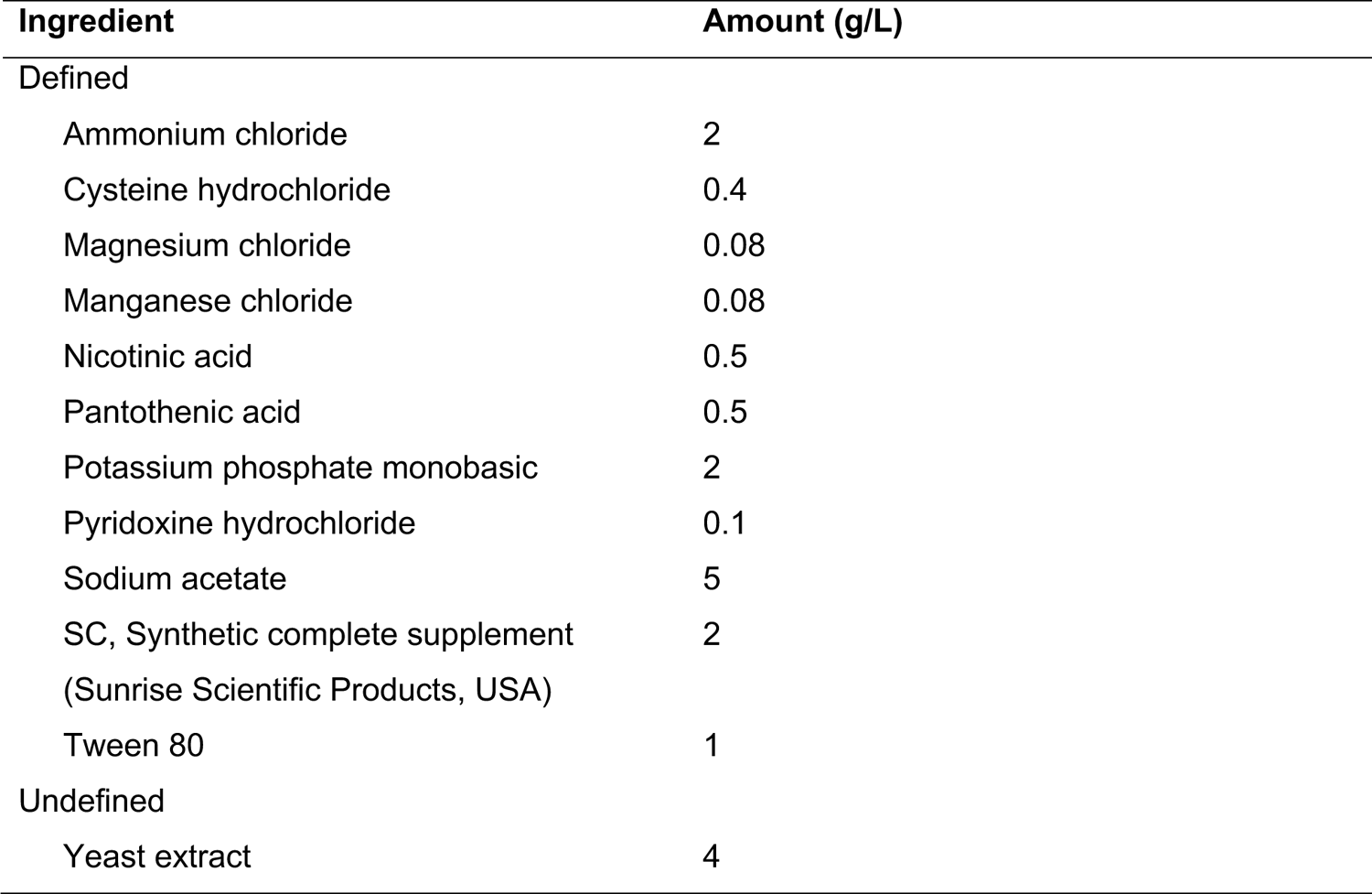
Composition of a semi-defined medium (SDM) recipe used to culture *Bifidobacterium* and *Lactobacillus* strains. Specific carbon sources (amygdalin and/or glucose) were added according to the experiments. Recipe was adapted from (1).

**Table S3.** Composition of a minimum medium (MM, pH 6.8) recipe used to culture transformed *E. coli* strains. Specific carbon sources (amygdalin, prunasin or glucose) were added according to the experiments. Recipe was adapted from (2).

| **Ingredient** | **Amount (g/L)** |
| --- | --- |
| Ammonium iron (III) citrate | 0.1 |
| Ammonium phosphate tetrahydrate | 4 |
| Boric acid | 0.003 |
| Citric acid | 1.55 |
| Cobalt (II) chloride hexahydrate | 0.003 |
| Copper (II) chloride dihydrate | 0.002 |
| Magnesium sulfate | 0.59 |
| Manganese (II) chloride tetrahydrate | 0.015 |
| Potassium phosphate monobasic | 13.3 |
| Sodium molybdate dihydrate | 0.002 |
| Zinc sulfate heptahydrate | 0.034 |

## Notes

### Competing Interest Statement

The authors have declared no competing interest.

